# Tandemly repeated genes promote RNAi-mediated heterochromatin formation via an anti-silencing factor Epe1 in fission yeast

**DOI:** 10.1101/2022.07.04.498379

**Authors:** Takahiro Asanuma, Soichi Inagaki, Tetsuji Kakutani, Hiroyuki Aburatani, Yota Murakami

## Abstract

In most eukaryotes, constitutive heterochromatin, defined by histone H3 lysine 9 methylation (H3K9me), is enriched on repetitive DNA, such as pericentromeric repeats and transposons. Furthermore, repetitive transgenes also induce heterochromatin formation in diverse model organisms. However, the mechanisms that promote heterochromatin formation at repetitive DNA elements are still not clear. Here, using fission yeast, we show that tandemly repeated mRNA genes promote RNA interference (RNAi)-mediated heterochromatin formation in co-operation with an anti-silencing factor, Epe1. Although the presence of tandemly repeated genes itself does not cause heterochromatin formation, once complementary small RNAs are artificially supplied in *trans*, the RNAi machinery assembled on the repeated genes starts producing cognate small RNAs in *cis* to autonomously maintain heterochromatin at these sites. The establishment of this “repeat-induced RNAi” depends on the copy number of repeated genes and also requires Epe1, which is known to remove H3K9me and derepress the transcription of genes underlying heterochromatin. Analogous to repeated genes, the DNA sequence underlying constitutive heterochromatin encodes widespread transcription start sites (TSSs), from which Epe1 activates ncRNA transcription to promote RNAi-mediated heterochromatin formation. Our results suggest that, when repetitive transcription units underlie heterochromatin, Epe1 generates sufficient transcripts for the activation of RNAi without disruption of heterochromatin.

## Introduction

In eukaryotic cells, histone H3 lysine 9 di- or trimethylation (H3K9me) defines heterochromatin, which represses the transcription and recombination of underlying DNA sequences. Consistent with its silent nature, constitutive heterochromatin is enriched on repetitive DNA elements, such as pericentromeric repeats and transposons, and maintains genome stability. However, the DNA sequences of such repetitive elements are not conserved across species (Saksouk et al. 2015). Furthermore, it has been reported that transformation of foreign DNA into higher eukaryotes often gives rise to repetitive transgenes, which leads to heterochromatin formation. Because the silencing of these repetitive transgenes depends on transgene copy number, this phenomenon is often referred to as repeat-induced gene silencing (RIGS) (Henikoff 1998). These facts suggest that it is the repetition itself that promotes heterochromatin formation; however, the mechanisms for this universal phenomenon are still not clear (Wang et al. 2006; Gladyshev and Kleckner 2017; Pal-Bhadra et al. 2004; Luo and Chen 2007; Schubert et al. 2004).

In fission yeast, the RNAi pathway promotes heterochromatin formation at pericentromeric and subtelomeric regions, and at the mating-type locus (Cam et al. 2005). These constitutive heterochromatin regions are defined by the presence of H3K9me and consist of common sequences, namely, *dg*/*dh* elements. From *dg*/*dh* elements, non-coding RNAs (ncRNAs) are transcribed by RNA polymerase II (Pol2) to act as scaffolds for the assembly of the RNAi machinery (Kato et al. 2005; Djupedal et al. 2005). The RNA-induced transcriptional silencing (RITS) complex, which contains Argonaute protein, Ago1, recognizes nascent *dg*/*dh* ncRNAs via cognate small interfering RNAs (siRNAs) (Verdel et al. 2004). This results in recruitment of the CLRC complex, which contains the Suv39h homolog, Clr4 that directs all H3K9me in this organism (Bayne et al. 2010). The RITS complex also recruits the RNA-dependent RNA polymerase complex (RDRC) to nascent ncRNAs (Motamedi et al. 2004). The RDRC synthesizes double-stranded RNA (dsRNA), which is processed by Dicer (Dcr1) to siRNAs. These secondary siRNAs function in the further recruitment of the RITS complex, thereby forming a positive feedback loop of siRNA production (Colmenares et al. 2007). Furthermore, the presence of H3K9me itself promotes the localization of the RITS complex and the RDRC (Noma et al. 2004; Hayashi et al. 2012; Rougemaille et al. 2012), indicating that heterochromatin formation and the RNAi pathway mutually reinforce each other. However, contrary to these findings, previous studies revealed that, even when cognate siRNA is artificially supplied in *trans*, the RNAi pathway hardly induces heterochromatin formation at euchromatic mRNA genes (Kowalik et al. 2015; Simmer et al. 2010; Iida et al. 2008). Furthermore, even if heterochromatin is formed at these sites, the RNAi pathway cannot maintain heterochromatin without cognate siRNA supplied in *trans* (Iida et al. 2008; Yu et al. 2018). Thus, these results raise the question of how RNAi-mediated heterochromatin formation is restricted to *dg/dh* ncRNAs and does not function on other mRNAs.

Recent studies using artificial tethering of Clr4 at euchromatin have shown that ectopically deposited H3K9me is actively removed, meaning that the epigenetic inheritance of H3K9me is strictly limited in the cell (Audergon et al. 2015; Ragunathan et al. 2015). They demonstrated that the JmjC domain demethylase family protein, Epe1, is responsible for H3K9me demethylation in this organism. In the presence of Epe1, ectopic heterochromatin induced by tethering of Clr4 at euchromatin is rapidly erased following release of tethered Clr4; however, in the absence of Epe1, ectopic heterochromatin induced at the tethering site is maintained during mitosis and meiosis, even after the release of tethered Clr4. This self-propagation of heterochromatin occurs because Clr4 can bind to established H3K9me-heterochromatin via its chromodomain and promotes the spread of H3K9me to neighboring nucleosomes (Audergon et al. 2015; Ragunathan et al. 2015). Notably, despite its function as a heterochromatin eraser, Epe1 is recruited to constitutive heterochromatin through its interaction with the H3K9me binding protein, Swi6/HP1(Zofall and Grewal 2006). While previous studies have shown that Epe1 plays a role in the establishment of the boundary between constitutive heterochromatin and euchromatin (Ayoub et al. 2003; Wang et al. 2013), the fact that loss of Epe1 (*epe1Δ*) also results in fewer siRNAs and defective silencing at constitutive heterochromatin (Trewick et al. 2007) implies that Epe1 has another function, involving the RNAi pathway, in maintenance of heterochromatin integrity. However, its exact role is still not understood.

Here, we show that Epe1 induces ncRNA expression from *dg*/*dh* elements to facilitate the assembly of the RNAi machinery on heterochromatin. Notably, ncRNAs whose transcription was induced by Epe1 have widespread transcription start sites (TSSs). This result prompted us to hypothesize that multiple transcription units underlying constitutive heterochromatin enable Epe1 to supply sufficient RNA templates to allow activation of RNAi even under silent heterochromatin. Consistent with this hypothesis, we found that, like *dg*/*dh* elements, tandemly repeated mRNA genes can establish constitutive heterochromatin via the RNAi pathway. These results indicate that RNAi-mediated heterochromatin formation is promoted by repeated transcription units, regardless of whether they encode mRNAs or ncRNAs. Thus, although Epe1 primarily removes heterochromatin, it also promotes heterochromatin formation via the RNAi pathway when repeated genes underlie silent heterochromatin. Such dual and opposing roles of an anti-silencing factor provide a novel insight into how repetitive DNA elements are linked to constitutive heterochromatin.

## Results

### Epe1 promotes assembly of the RNAi machinery on constitutive heterochromatin

Activation of the RNAi pathway involves recognition of nascent target RNAs by the RITS complex and concomitant recruitment of the CLRC and the RDRC, and is thus accompanied by the assembly of these factors onto the chromatin of the corresponding region. To define the link between Epe1 and the RNAi pathway at *dg*/*dh* elements, we first examined the effect of Epe1 on the assembly of the RNAi machinery on constitutive heterochromatin. While loss of Epe1 resulted in dissociation of the RNAi machinery from pericentromeric heterochromatin (Fig. 1A; Supplemental Fig. S1A-C), overproduction of Epe1 (Epe1 OP) (Supplemental Fig. S1D-G) significantly promoted its assembly, with concomitant upregulation of siRNAs (Fig. 1A, C). Similar results were also obtained for other constitutive heterochromatin regions such as the mating-type locus (Supplemental Fig. S1H). Consistent with previous reports that Epe1 OP derepresses expression of a reporter gene inserted within constitutive heterochromatin (Zofall and Grewal 2006; Ayoub et al. 2003; Trewick et al. 2007) (Supplemental Fig. S1G), ncRNA expression and Pol2 occupancy at *dg*/*dh* elements were significantly increased by Epe1 OP (Fig. 1D; Supplemental Fig. S1I) (Bao et al. 2018). These results suggest that Epe1 promotes assembly of the RNAi machinery at constitutive heterochromatin by expressing *dg*/*dh* ncRNAs. Despite the high expression of ncRNAs in Epe1 OP cells, H3K9me levels at constitutive heterochromatin were maintained, probably because the hyper-activated RNAi replenished H3K9me (Fig. 1A-C; Supplemental Fig. S1H).

**Figure 1.**
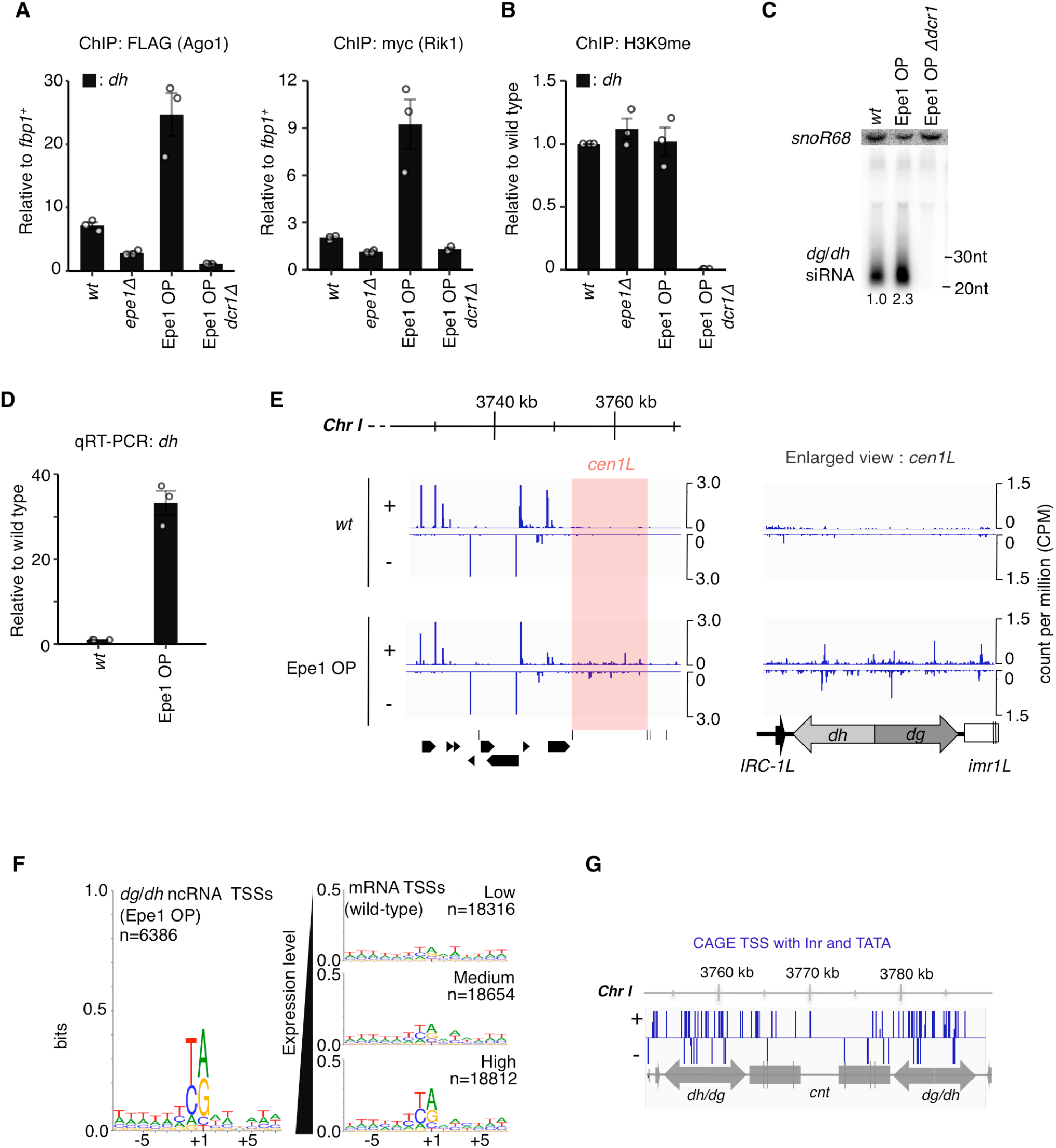
Epe1 overproduction promotes assembly of the RNAi machinery by inducing ncRNA expression from TSSs that are widespread across constitutive heterochromatin. (*A*) ChIP-qPCR of Ago1, a CLRC component, Rik1, and (*B*) H3K9me at pericentromeric *dh*. Error bars represent SEM; n = 3 biological replicates. (*C*) Northern blotting of *dg*/*dh* siRNAs. snoRNA58 (snoR58), loading control. Average signal intensities calculated from three independent experiments are shown. (*D*) qRT-PCR of *dh* transcripts relative to the wild type (*wt*). *(E)* CAGE-seq reads of *wt* or Epe1 OP in the vicinity of chromosome I left pericentromere (cen1L) and an enlarged view of cen1L are shown in a strand-specific manner (±). (*F*) The consensus sequence of TSSs in *dg*/*dh* elements with Epe1 OP (>0.05 CPM) was compared with those of mRNAs in the *wt*. TSSs of mRNAs were categorized into three groups according to expression strength. The number of unique TSSs in each category is indicated as “n.” (*G*) CAGE reads that have both Inr elements and a TATA-like A/T rich region were extracted and mapped to the chromosome I pericentromere.

### Epe1 induces ncRNA expression from transcription start sites that are widespread across constitutive heterochromatin

In wild-type cells, *dg*/*dh* ncRNAs are hardly detectable due to their transcriptional silencing and processing by the RNAi machinery. Therefore, the higher expression of *dg*/*dh* ncRNA in the Epe1 OP strain facilitated analysis of these ncRNAs. Cap Analysis of Gene Expression (CAGE)-seq, which can profile TSSs of capped RNAs transcribed by Pol2, revealed that Epe1 OP activates weak transcription from TSSs that are widespread across constitutive heterochromatin (Fig. 1E; Supplemental Fig. S2A-C). Consensus sequence analysis of these weak TSSs revealed that many of them have features in common with mRNA TSSs. Namely, the Y/R (Y = pyrimidine, R = purine) dinucleotide is enriched at -1/+1 positions (Fig. 1F), and an A/T rich region is observed -25–32 nt upstream (Supplemental Fig. S2D). These motifs correspond to an initiator (Inr) element and a TATA box of euchromatic mRNA genes, respectively (Li et al. 2015). Thus, these results revealed that *dg*/*dh* elements encode a distinctive structure with widespread TSSs, which have a preference for a core promoter structure identical to that of mRNAs (Fig. 1G).

To analyze the structure of *dg*/*dh* elements in more detail, we next performed 5’/3’ RACE to identify the TSSs and transcription termination sites (TTSs) of *dg*/*dh* ncRNAs. We chose *dg*/*dh* elements at the mating-type locus, namely, *cenH*, as a model for *dg*/*dh* elements in this study because, although *cenH* is highly homologous to pericentromeric *dg*/*dh* elements, it has a specific insertion that enables us to distinguish it from other *dg*/*dh* elements (Supplemental Fig. S3A). Furthermore, in addition to the RNAi pathway, the DNA-binding proteins, Atf1/Pcr1, also recruit Clr4 to maintain heterochromatin at the mating-type locus (Jia et al. 2004). This enables us to analyze transcripts induced by Epe1 in RNAi-defective mutants without no heterochromatin disruption.

In the absence of heterochromatin (Supplemental Fig. S3D, *clr4Δ*), convergent major TSSs were detected on *cenH*, while minor widespread TSSs sandwiched between these major sites were also detected on both strands. This result is consistent with a previous report that multiple TSSs were detected on both strands of pericentromeric *dg*/*dh* fragments in *clr4Δ* cells using 5’RACE analysis (Buscaino et al. 2013). In wild-type cells, it was impossible to clone 5’RACE products of the *cenH* ncRNA due to its extremely low expression levels. Therefore, to identify TSSs activated in the presence of heterochromatin, we used RNAi-defective *dcr1Δ* cells. In RNAi-defective mutants, ncRNA expression was detectable even in the presence of heterochromatin because they are not processed by the RNAi pathway (Supplemental Fig. S3B). Notably, in the presence of heterochromatin, minor widespread TSSs were activated (Supplemental Fig. S3D, *dcr1Δ*), resulting in smearing of 5’RACE products when analyzed by electrophoresis (Supplemental Fig. S3B, *dcr1Δ*). This activation of widespread TSSs is dependent on Epe1 because additional deletion of *epe1*^*+*^ dramatically abrogated their activation (Supplemental Fig. S3D, *dcr1Δepe1Δ*), indicated by the reappearance of a convergent band in electrophoresis (Supplemental Fig. S3B, *dcr1Δepe1Δ*). A a single deletion of *epe1*^*+*^ also resulted in repression of widespread TSSs, with activation of only major TSSs (Supplemental Fig. S3D, *epe1Δ*). The residual activation of major TSSs explains the presence of ncRNA transcription in *epe1Δ* cells (Supplemental Fig. S3B), which was also reported in a previous study (Zofall and Grewal 2006). By contrast, consistent with results obtained using CAGE-seq, Epe1 OP caused hyper-activation of the widespread TSSs (Supplemental Fig. S3D, Epe1 OP), which again appeared as smearing products in electrophoresis (Supplemental Fig. S3B, Epe1 OP). Thus, these results indicate that the *dg*/*dh* element consists of two types of TSS: Epe1-independent TSSs, which are resistant to silencing by heterochromatin, and widespread TSSs, whose expression is dependent on Epe1 in the presence of heterochromatin.

As mentioned above, Epe1 OP causes derepression of a reporter gene inserted within constitutive heterochromatin. To investigate the nature of heterochromatic transcription induced by Epe1, we next performed 5’/3’RACE analysis on an *ura4*^*+*^ gene integrated into *cenH* (*Kint2::ura4*^*+*^). The expression of *Kint2::ura4*^*+*^ is derepressed by Epe1 OP in the presence of heterochromatin (Supplemental Fig. S1G,S3F). Our 5’/3’RACE analysis revealed that Epe1 promotes expression of *Kint2::ura4*^*+*^via TSSs/TTSs that are almost identical to endogenous ones (Supplemental Fig. S3C,E), indicating that Epe1 induces transcription from heterochromatin by following the underlying DNA sequence. On the other hand, our 3’RACE analysis identified multiple TTSs in *cenH* ncRNA, as previously reported at pericentromeric *dg*/*dh* elements using PolyA-seq (Yu et al. 2014). Compared with its effect on TSSs, the presence or absence of Epe1 did not significantly affect the distribution of TTSs (Supplemental Fig. S3D), leading us to speculate that the function of Epe1 in the RNAi pathway is exerted by activation of these widespread TSSs in *dg*/*dh* elements.

### A fragment containing only widespread TSSs can establish heterochromatin

Our 5’RACE analysis revealed that *dg*/*dh* elements at the mating-type locus (*cenH*) include two types of TSS: Epe1-independent/silencing-resistant convergent TSSs and Epe1-dependent widespread TSSs. To determine which TSS element is required for RNAi-mediated heterochromatin formation, a series of truncated *cenH* fragments were cloned into plasmid-based minichromosomes (Buscaino et al. 2013) and transformed into the *h*^-S^ strain, in which the native *cenH* region at the mating-type locus is completely lost. This enabled specific evaluation of siRNA production and H3K9me accumulation at the truncated *cenH* fragment. This analysis revealed that a 1.5 kb fragment containing only widespread TSSs can establish heterochromatin, which is accompanied by production of siRNAs from this fragment (Supplemental Fig. S4). Thus, it appears that Epe1-independent TSSs, namely, silencing-resistant TSSs, are not essential for RNAi-mediated heterochromatin formation.

To examine the contribution of widespread TSSs to RNAi-mediated heterochromatin formation, it would be necessary to destroy widespread TSSs across the whole 1.5 kb fragment. However, such an experiment was not feasible because the high number of TSSs identified in this fragment by 5’ RACE (the distance between the closest neighboring TSSs was only 3 bp; Supplemental Data S1) meant that the base substitutions necessary to remove these TSSs would result in a DNA sequence that is significantly different from the original.

### Repeated mRNA genes promote RNAi-mediated heterochromatin formation more efficiently than a single gene

A paradox of RNAi-mediated heterochromatin formation is that “silent” heterochromatin formation requires “transcription” of the corresponding region. Therefore, the presence of widespread TSSs in *dg/dh* elements led us to speculate that these multiple TSSs enable Epe1 to supply sufficient RNA template for RNAi to occur, even under silent heterochromatin. Previous studies using hairpin RNA demonstrate that targeting the RNAi to mRNAs with artificial siRNAs in *trans* (*trans*-acting RNAi) hardly induces *de novo* heterochromatin formation (Kowalik et al. 2015; Simmer et al. 2010; Iida et al. 2008), and it has been proposed that euchromatic mRNA genes are protected from siRNA-directed heterochromatin formation by unknown mechanisms. We therefore hypothesized that when target mRNA genes are repeated at a single genomic locus, sufficient RNA templates are supplied for the RNAi to occur even in the presence of heterochromatin, as occurs in *dg*/*dh* elements with multiple TSSs.

To test this hypothesis, we first generated a series of strains in which the reporter gene *ade6*^*+*^ was tandemly repeated in increasing copy number, ranging from one to eight, at the endogenous *ura4*^*+*^ locus (Fig. 2A and Supplemental Fig. S5A-C). Northern blot analysis and RT-PCR with a strand-specific primer detected no obvious readthrough transcript or an antisense transcript specific for the *ade6*^*+*^ repeats (Supplemental Fig. S5D,E). Furthermore, qRT-PCR with an oligodT primer confirmed that the expression levels of the *ade6*^*+*^ repeats are consistent with copy number (Supplemental Fig. S5F). We chose the *ade6*^*+*^ gene as a reporter because silencing of *ade6*^*+*^ results in red-pink colony formation on indicator plates. Silencing assays showed that both the maximum copy number *ade6*^*+*^x8 strain and the minimum copy number *ade6*^*+*^x1 strain formed only white (*ade6*-expressing) colonies (Fig. 2B,C), suggesting that repetition itself does not cause silencing of *ade6*^*+*^ in this organism. We next created two hairpin RNA constructs, *ade6*-hp I and II, which produce small RNAs complementary to a portion of the *ade6* ORF; the *ade6*-hp I and *ade6*-hp II constructs have a hairpin structure of 250 bp and 750 bp, respectively (Supplemental Fig. S6). These constructs were integrated into the endogenous *leu1*^*+*^ locus to induce *trans*-acting RNAi (Fig. 2A). Because post-transcriptional gene silencing is negligible in fission yeast (Kowalik et al. 2015; Simmer et al. 2010; Iida et al. 2008), emergence of the red-pink (*ade6*-repressed) phenotype with *trans*-acting RNAi reflects heterochromatin formation on *ade6*^*+*^. Consistent with previous studies, when *ade6*-hp I or II were expressed in the *ade6*^*+*^x1 strains, red-pink (*ade6*-repressed) colonies emerged from white (*ade6*-expressing) colonies at very low rates of 0% and 0.2%, respectively (Fig. 2C). On the other hand, when *ade6*-hp I or II were expressed in the *ade6*^*+*^x8 strains, the emergence of red-pink (*ade6*-repressed) colonies significantly increased to 0.2% and 10.7%, respectively (Fig. 2C). The variation in red-pink (*ade6*-repressed) colony formation between *ade6*-hp II and *ade6*-hp I expressing cells is probably due to the difference in hairpin lengths. On the other hand, the stability of the red-pink (*ade6*-repressed) phenotype in the *ade6*^*+*^x8 strain, which was evaluated by the frequency with which the phenotype switched from red-pink (*ade6*-repressed) to white (*ade6*-expressing), was very high with both hairpin RNA constructs; more than 97% of cells maintained the red-pink (*ade6*-repressed) phenotype (Fig. 2C). By contrast, the red-pink (*ade6*-repressed) phenotype established with the *ade6*^*+*^x1 strain using *ade6*-hp II was very unstable; more than 90% of red-pink (*ade6*-repressed) cells reverted to the white (*ade6*-expressing) phenotype (Fig. 2C). Thus, these results suggest that repeated mRNA genes are a more efficient target of *trans*-acting RNAi than a single gene, mainly because maintenance of established heterochromatin is improved with repeated genes.

**Figure 2.**
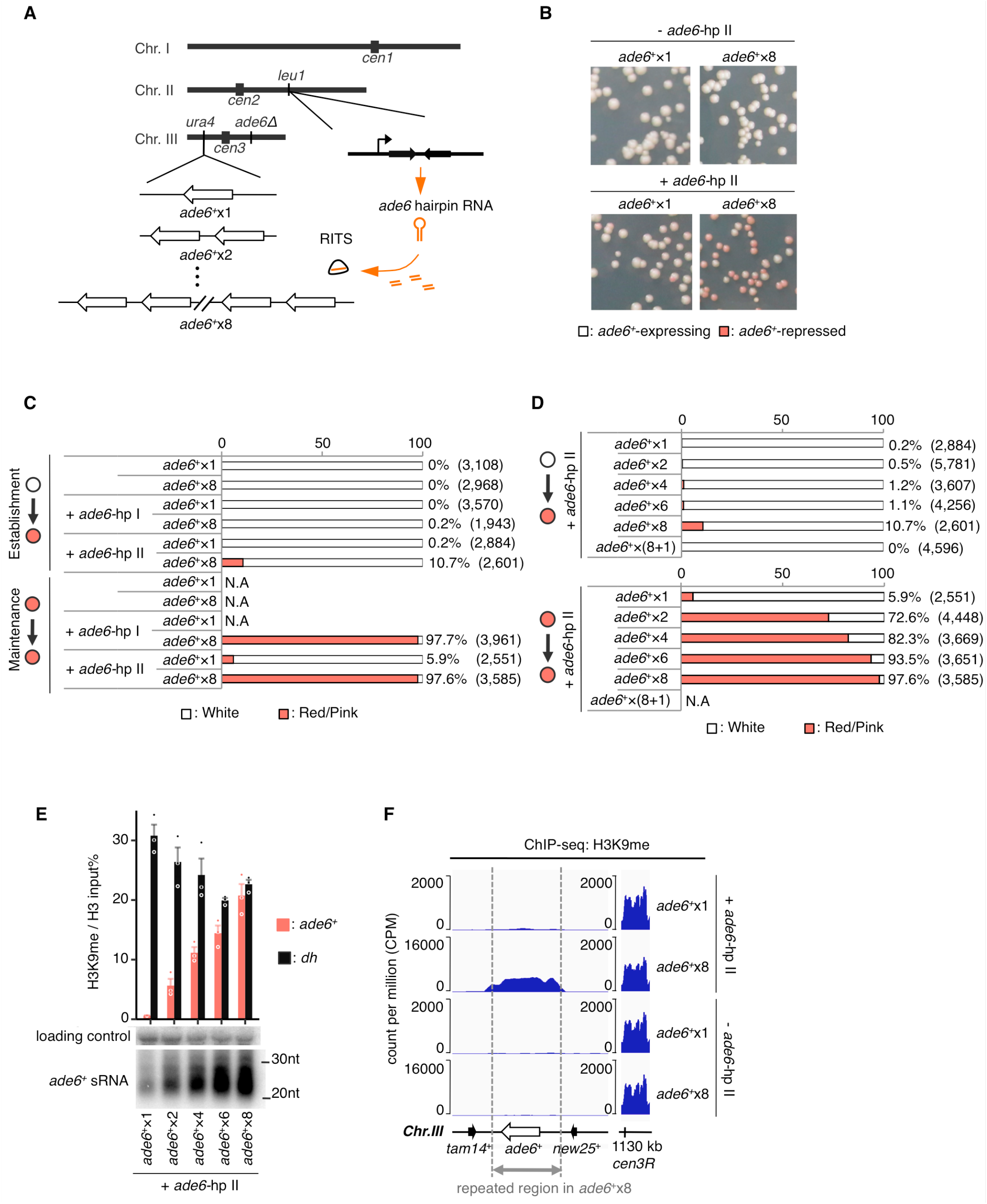
Repeated genes promote RNAi-mediated heterochromatin formation. (*A*) Diagram of experimental scheme for artificial targeting of RNAi to the repeated reporter gene, *ade6*^*+*^. (*B*) Representative images of silencing assay for *ade6*^+^x1 and *ade6*^+^x8 strains with or without *ade6*-hp II. (*C*)(*D*) Using silencing assay, the efficiency of establishment or maintenance of the *ade6*-repressed state was evaluated. The number of scored colonies is shown in parentheses. (*C*) *ade6*^*+*^x1 and *ade6*^*+*^x8 strains without/with hairpin RNAs (*ade6*-hp I and II). (*D*) A series of *ade6*^*+*^ repeat strains with *ade6*-hp II. (*E*) Upper, ChIP-qPCR of H3K9me with the indicated strains after red-pink (*ade6*-repressed) clones were selected. Error bars represent SEM; n = 3 biological replicates. Lower, Northern blot of *ade6*^*+*^ siRNA using the same cells. A non-specific band was used as a loading control. (*F*) ChIP-seq analysis of H3K9me with cells used in (*E*) and negative controls. To facilitate visualization, reads were mapped on *ade6*^*+*^x1 construct. Results of *ade6*^*+*^x8 strains are scaled by a factor of 8 relative to *ade6*^*+*^x1.

To further examine the effect of repetition on heterochromatin formation via the RNAi pathway, we evaluated the establishment and maintenance of the red-pink (*ade6*-repressed) phenotype using strains in which a series of *ade6*^*+*^ repeats was combined with *ade6*-hp II (Fig. 2D). As expected, an increase in the number of repeats tended to promote heterochromatin formation, and this trend was more pronounced with respect to its maintenance (Fig. 2D). Consistent with the results of this silencing assay, when levels of H3K9me on *ade6*^*+*^ were evaluated using cells isolated from red-pink (*ade6*-repressed) clones, a clear correlation between H3K9me levels on *ade6*^*+*^ and copy number was observed. In the maximum *ade6*^*+*^x8 strain, the level of H3K9me on *ade6*^*+*^ was the same as that on pericentromeric heterochromatin (Fig. 2E, F).

### The promotion of RNAi-mediated heterochromatin formation by repeated mRNA genes is not due to their gene dosage

To determine whether the effect of repeated mRNA genes on RNAi-mediated heterochromatin formation depends on its gene dosage, we next performed the silencing assay using an *ade6*^*+*^x(8+1) strain, in which the endogenous *ade6*^*+*^ gene coexists at a distance from the *ade6*^*+*^x8 allele (Supplemental Fig. S7A, left panel). In contrast to the *ade6*^*+*^x8 strain, *trans*-acting RNAi using *ade6*-hp II failed to induce formation of red-pink (*ade6*-repressed) colonies in the *ade6*^*+*^x(8+1) strain (Fig. 2D). In principle, when multiple *ade6*^*+*^ genes exist in a cell, expression of just one *ade6*^*+*^ gene will result in the white (*ade6*-expressing) phenotype, even if the rest of *ade6*^*+*^ genes are silenced. Therefore, the absence of red-pink (*ade6*-repressed) colony formation in the *ade6*^*+*^x(8+1) strain suggests that there may be a bias in the ease of heterochromatin formation between the *ade6*^*+*^x8 and isolated the *ade6*^*+*^ gene. However, it is difficult to examine this possibility with this strain because its white (*ade6*-expressing) phenotype made it difficult to identify a clone in which heterochromatin is established on *ade6*^*+*^ genes. To address this, we next combined the endogenous *ade6*^*+*^ gene with the *ade6*^*+*^x8 allele, which is already heterochromatinized by *trans*-acting RNAi with *ade6*-hp II (Supplemental Fig. S7A, right panel). The resultant *ade6*^*+*^x(8*+1) strain with *ade6*-hp II formed red-pink (*ade6*-repressed) colonies stochastically; however, these colonies failed to stably maintain their red-pink (*ade6*-repressed) phenotype, in contrast to the cognate *ade6*^*+*^x8 strain with *ade6*-hp II (Supplemental Fig. S7B). This result indicates that RNAi inefficiently targets an isolated *ade6*^*+*^ gene, even in the presence of repeated *ade6*^*+*^ genes. Consistently, semiquantitative ChIP-PCR, which can discriminate between *ade6*^*+*^x8 and an isolated *ade6*^*+*^, showed that H3K9me was hardly deposited on an isolated *ade6*^*+*^ gene in this strain (Supplemental Fig. S7C, D). Thus, these results indicate that the promotion of RNAi-mediated heterochromatin formation by repeated mRNA genes is not due to their gene dosage.

### Repeated mRNA genes promote *cis*-acting RNAi

To explore in detail why repeated genes promote RNAi-mediated heterochromatin formation, we next examined the effect of repeated genes on siRNA production. Notably, we found that heterochromatin formation at the *ade6*^*+*^x8 allele was accompanied by significant production of novel siRNAs that were not encoded by *ade6*-hp II, i.e., secondary siRNAs (Fig. 3A). This result indicated that, with the *ade6*^*+*^x8 allele, *trans*-acting RNAi efficiently activates the RNAi pathway in *cis* on *ade6*^*+*^ mRNAs (*cis*-acting RNAi) (Fig. 3B). On the other hand, although secondary siRNA was also detected with the *ade6*^*+*^x1 allele, as previously reported (Simmer et al. 2010), it was present in only very small amounts compared with those observed for the *ade6*^*+*^x8 allele (Fig. 3A). This result indicates that repeated mRNA genes allow *trans*-acting RNAi to activate the *cis*-acting RNAi more efficiently than a single gene. Consistently, Northern blot analysis using a series of *ade6*^*+*^ repeat strains showed that total *ade6* siRNA levels, which include both primary siRNAs from *ade6*-hp II and secondary siRNAs from *cis*-acting RNAi, correlated with the copy number of *ade6*^*+*^ repeats (Fig. 2E). Thus, the activation of *cis*-acting RNAi explains accumulation of H3K9me, which was correlated with copy number of target genes.

**Figure 3.**
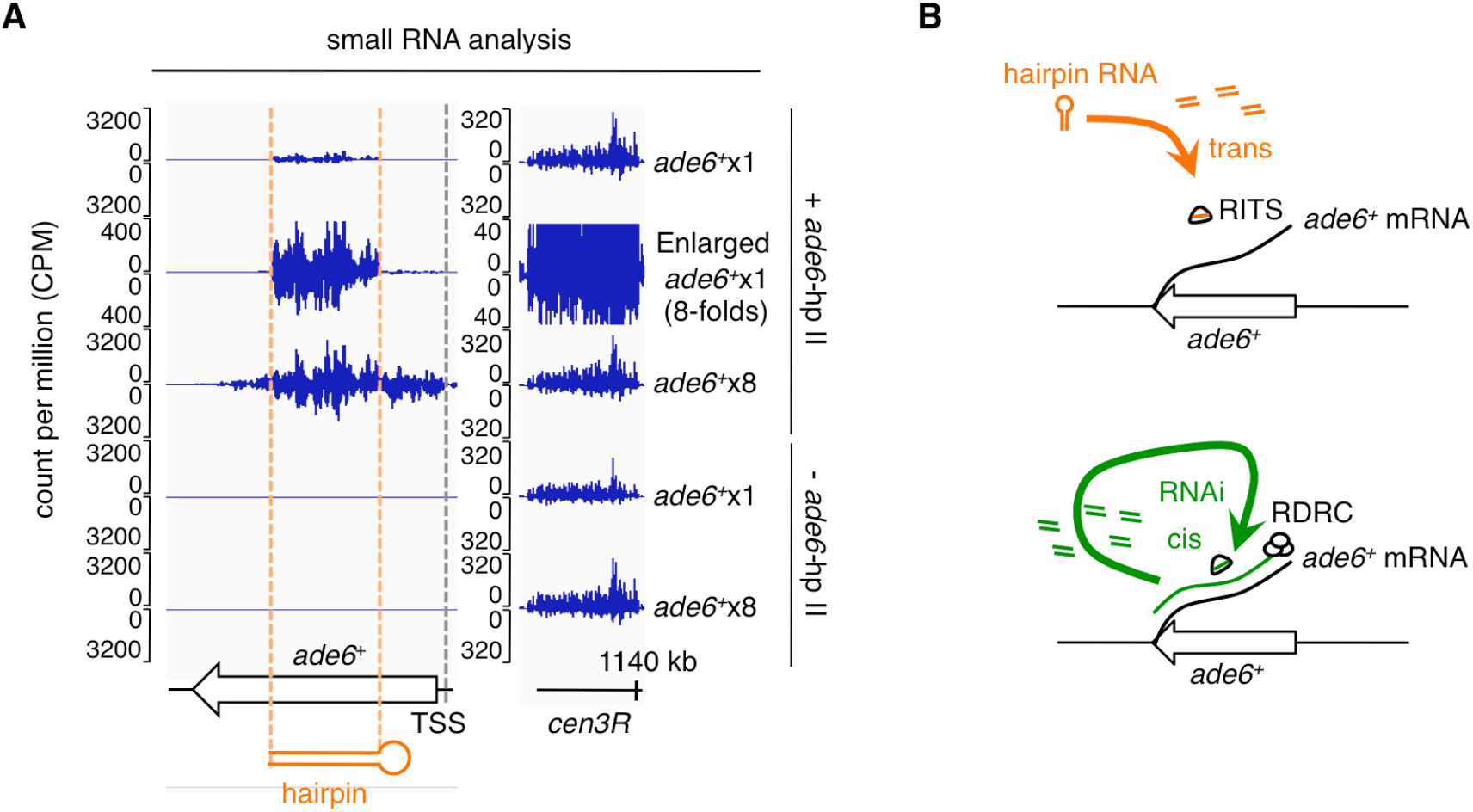
Repeated genes promote *cis*-acting RNAi more efficiently than a single gene. (*A*) Small RNA reads mapping to *ade6*^*+*^ and part of the chromosome III pericentromere are shown. Dashed orange lines mark a region targeted by *trans*-acting RNAi (*ade6*-hp II). An 8-fold enlarged view of *ade6*^*+*^x1 is also shown to take into account the effect of copy number. For other small RNAs mapped around *ade6*^*+*^, see also Supplemental Fig. S11. (*B*) Schematic diagram of the RNAi pathway targeted to *ade6*^*+*^ mRNAs by two different routes. Upper, *trans*-acting RNAi, in which siRNAs derived from *ade6* hairpin RNAs direct the RNAi pathway to *ade6*^*+*^ mRNAs in *trans*. Lower, *cis*-acting RNAi, in which siRNAs derived from *ade6*^*+*^ mRNAs direct the RNAi pathway to *ade6*^*+*^ mRNAs in *cis*. When *trans*-acting RNAi activates *cis*-acting RNAi, dsRNA synthesis by the RDRC and subsequent processing by Dicer produces secondary siRNAs, which are not encoded by hairpin RNAs.

### Repeated mRNA genes can establish autonomous *cis*-acting RNAi (Repeat-induced RNAi)

Previous studies using hairpin RNA demonstrated that, even if *trans*-acting RNAi succeeds in *de novo* heterochromatin formation at euchromatic mRNA genes, established ectopic heterochromatin is not maintained after removal of hairpin RNA (Iida et al. 2008; Yu et al. 2018). To test whether the continuous presence of *trans*-acting RNAi is necessary for maintenance of ectopic heterochromatin on the *ade6*^*+*^x8 allele, we segregated *ade6*-hp II from the *ade6*^*+*^x8 allele by crossing with cells that do not have the *ade6*-hp II allele (*ade6*-hp II*Δ*) (Fig. 4A). When combined with *ade6*-hp II*Δ*, the *ade6*^*+*^x8 cells continued to exhibit the same red-pink (*ade6*-repressed) phenotype as the original *ade6*^*+*^x8 cells combined with *ade6*-hp II (Fig. 4A), and H3K9me on *ade6*^*+*^x8 in the absence of *ade6*-hp II was indeed maintained at the same level as seen when *ade6*-hp II is present (Fig. 4B). Consistent with this, *ade6*^*+*^ siRNAs were autonomously produced from the *ade6*^*+*^x8 allele even after the removal of *ade6*-hp II (Fig. 4C). siRNAs derived from *ade6*^*+*^x8 exhibited the same properties as native *dg*/*dh* siRNAs (Supplemental Fig. S8), and resultant heterochromatin on *ade6*^*+*^x8 allele was inherited through both mitosis and meiosis in an RNAi components-dependent manner (Fig. 4C, D, and F). These results indicate that autonomous *cis*-acting RNAi is established on the *ade6*^*+*^x8 allele and maintains heterochromatin at this site. Thus, although repetition of *ade6*^*+*^ itself does not cause heterochromatin formation, once recognized, it starts to function as a platform for the RNAi pathway, in the same way as *dg*/*dh* elements. To the best of our knowledge, this is the first RIGS phenomenon observed in fission yeast. Since the term of RIGS (repeat-induced gene silencing) does not define its mechanism, we have therefore named this autonomous *cis*-acting RNAi “repeat-induced RNAi,” in analogy to RIGS.

**Figure 4.**
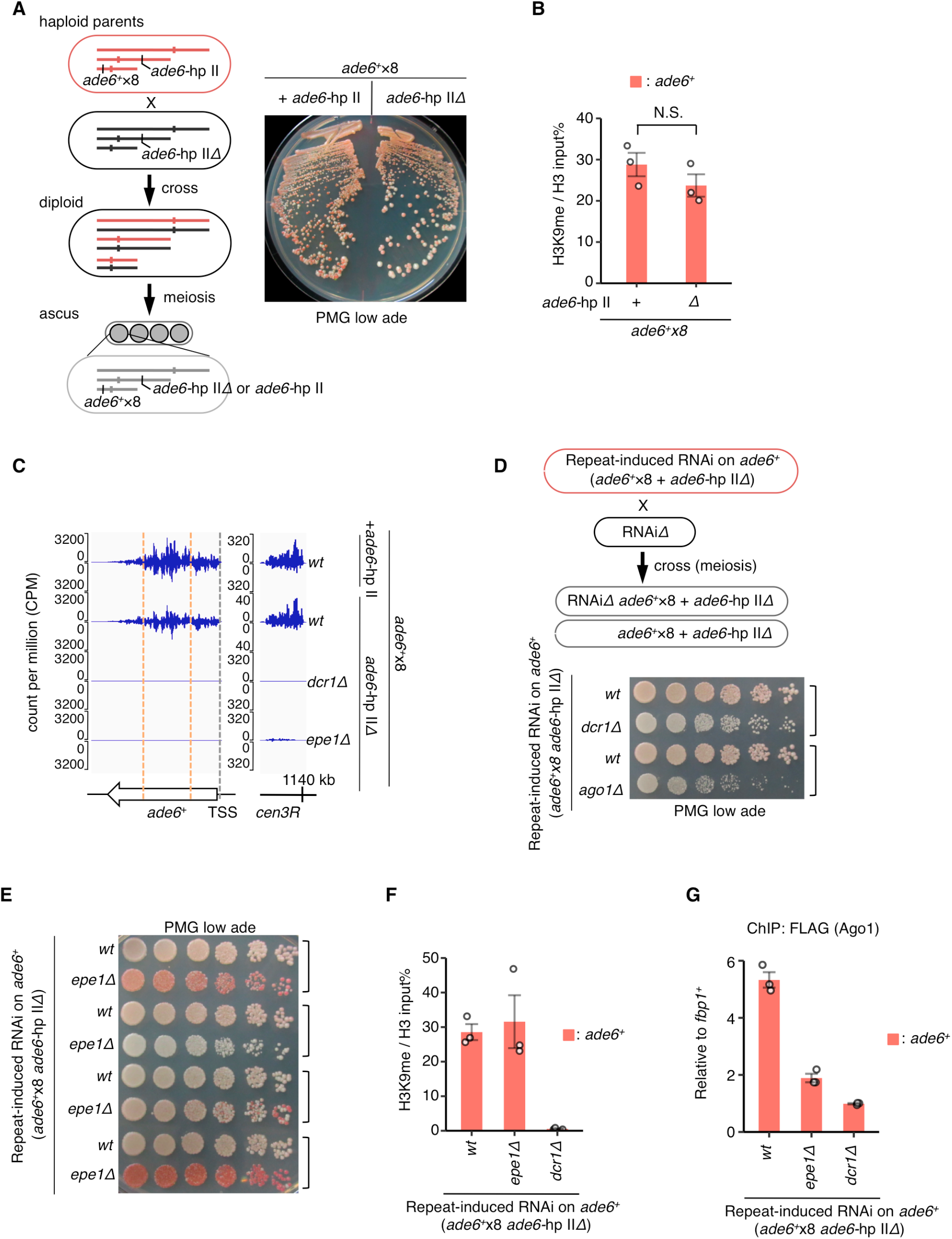
Epe1 is required for repeat-induced RNAi. (*A*) Silencing assay of *ade6*^*+*^x8 strains after removal of the hairpin RNA construct. (*B*) ChIP-qPCR of H3K9me with progenies that inherited cognate *ade6*^*+*^x8 alleles with/without *ade6*-hp II. (*C*) Small RNA-seq of the indicated strains with/without *ade6*-hp II. Note that the result of *wt ade6*^*+*^x8 with *ade6*-hp II is shared with Fig. 3. (*D*)(*E*) Silencing assay of *ade6*^*+*^x8 allele combined with RNAi-defective mutants, (*D*) *dcr1Δ, ago1Δ*, and (*E*) *epe1Δ* cells. Wild-type cells derived from the same ascus, which inherit the cognate *ade6*^*+*^x8 allele, are also indicated with close square brackets. (*F*)(*G*) ChIP-qPCR of (E) H3K9me and (*G*) Ago1 at the *ade6*^*+*^x8 allele with indicated strains. Error bars represent SEM; n = 3 biological replicates. N.S., not significant (P = 0.27, two-sided Student’s t-test).

### Establishment of repeat-induced RNAi depends on the copy number of repeated mRNA genes and requires an anti-silencing factor Epe1

Because *trans*-acting RNAi was likely to promote *cis*-acting RNAi in correlation with the copy number (Fig. 2E), we next examined whether the establishment of repeat-induced RNAi also depends on the copy number of target gene repeats. To examine this, we removed *ade6*-hp II from each *ade6*^*+*^x1, x2, x4, x6, x8 strain in which *trans*-acting RNAi established heterochromatin on *ade6*^*+*^ genes (Fig. 5A). Specifically, red-pink (*ade6*-repressed) cells of each *ade6*^*+*^ repeat strain were crossed with *ade6*-hp II*Δ* cells to segregate *ade6*-hp II, and the percentage of progenies that maintain the red-pink (*ade6*-repressed) phenotype without *ade6*-hp II was assessed. This revealed that most of the progenies derived from cells with four or more copies of the *ade6*^*+*^ gene showed the red-pink (*ade6*-repressed) phenotype, while those derived from *ade6*^+^x1 or x2 cells showed only the white (*ade6*-expressing) phenotype (Fig. 5A). Consistent with their phenotype, autonomous production of *ade6* siRNAs and heterochromatin maintenance were observed with the red-pink (*ade6*-repressed) progenies but not the white (*ade6*-expressing) progenies derived from *ade6*^+^x1 or x2 strains (Fig. 5B). Thus, these results indicate that the establishment of repeat-induced RNAi requires a minimal number of repeated mRNA genes.

**Figure 5.**
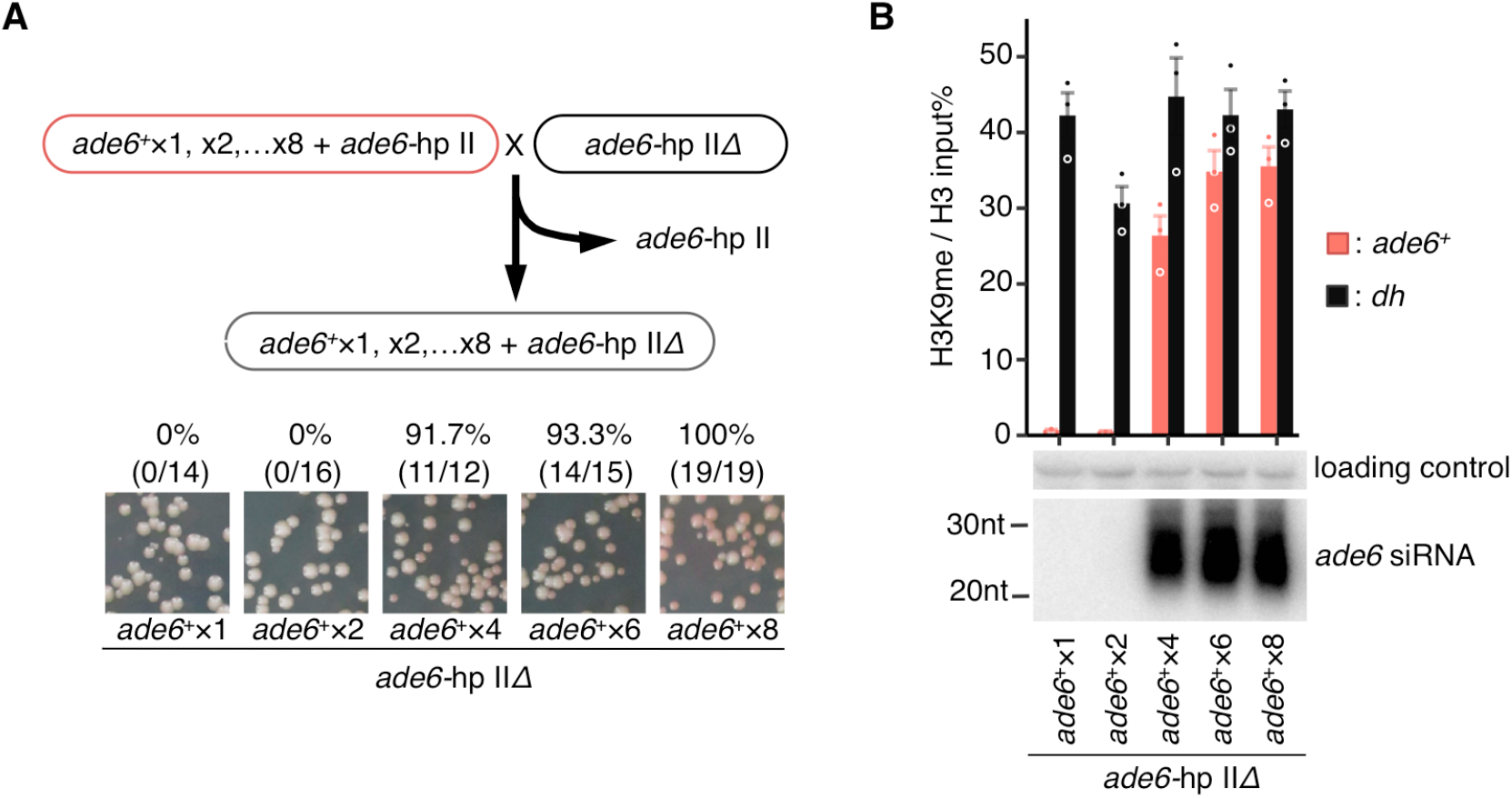
Autonomous *cis*-acting RNAi is not established with *ade6*^*+*^x1 or x2 alleles. (*A*) Red cells, in which a series of *ade6*^*+*^ repeat alleles were heterochromatinized by *ade6*-hp II, were crossed with *ade6*-hp II*Δ* cells to segregate the *ade6*^+^ repeat allele from the *ade6*-hp II. The percentage of progenies that formed red-pink (*ade6*-repressed) colonies without *ade6*-hp II is indicated. (*B*) Upper, ChIP-qPCR of H3K9me, and lower, Northern blot analysis for *ade6* siRNAs for the indicated strains. Non-specific bands produced by the *ade6* probe were used as a loading control.

We next examined whether Epe1 is also required for repeat-induced RNAi at *ade6*^*+*^x8 because Epe1 is required for the assembly of the RNAi machinery and siRNA production at *dg*/*dh* elements (Fig. 1). Consistent with the establishment of RNAi-mediated heterochromatin formation, the RITS complex was localized at the *ade6*^*+*^x8 locus, where repeat-induced RNAi maintains heterochromatin (Fig. 4G). Notably, loss of Epe1 significantly decreased its localization (Fig. 4G) and depleted *ade6* siRNAs, as it does in the RNA pathway at *dg*/*dh* elements (Fig. 4C). Thus, these results indicate that the establishment of repeat-induced RNAi not only depends on the copy number of repeated mRNA genes but also requires the anti-silencing factor, Epe1. Note that H3K9me on the *ade6*^*+*^x8 gene was maintained in *epe1Δ* cells with variegated silencing (Fig. 4E,F) because heterochromatin can persist by self-propagation in the absence of Epe1 (Audergon et al. 2015; Ragunathan et al. 2015; Trewick et al. 2007).

### Epe1 plays opposing roles in epigenetic inheritance of H3K9me in a repeat-dependent manner

Our results demonstrate that Epe1 promotes the assembly of RNAi machinery to deposit H3K9me at *dg*/*dh* elements, although Epe1 removes ectopically deposited H3K9me at euchromatin. To understand how Epe1 plays such opposing roles in epigenetic inheritance of H3K9me, we used our repeat-induced RNAi system. First, the role of Epe1 in repeat-induced RNAi was examined using *trans*-acting RNAi. Because loss of Epe1 does not abrogate production of *ade6* siRNAs from *ade6*-hp II (Supplemental Fig. S6), *trans*-acting RNAi enables evaluation of the effects of *epe1Δ* on siRNA-directed heterochromatin formation. In *epe1Δ* cells, *trans*-acting RNAi established heterochromatin more efficiently than in wild-type cells at both the *ade6*^*+*^x1 and the *ade6*^*+*^x8 allele (Supplemental Fig. S9), indicating that, although euchromatic mRNA genes are susceptible to siRNA-directed heterochromatin formation, Epe1 primarily suppresses it, regardless of whether or not the target gene is repeated. This idea was further supported by the fact that, once established, robust accumulation of H3K9me was observed at both the *ade6*^*+*^x1 and *ade6*^*+*^x8 alleles in *epe1Δ* cells (Fig. 6A). Notably, these robust ectopic heterochromatins were not accompanied by effective production of secondary siRNAs even at the *ade6*^*+*^x8 allele (Fig. 6B). This result indicates that the presence of H3K9me, which facilitates the recruitment of the RITS complex and the RDRC (Noma et al. 2004; Hayashi et al. 2012; Rougemaille et al. 2012), is insufficient to allow *trans*-acting RNAi to activate *cis*-acting RNAi, and that Epe1 must also be present, presumably to supply RNA templates for *cis*-acting RNAi.

**Figure 6.**
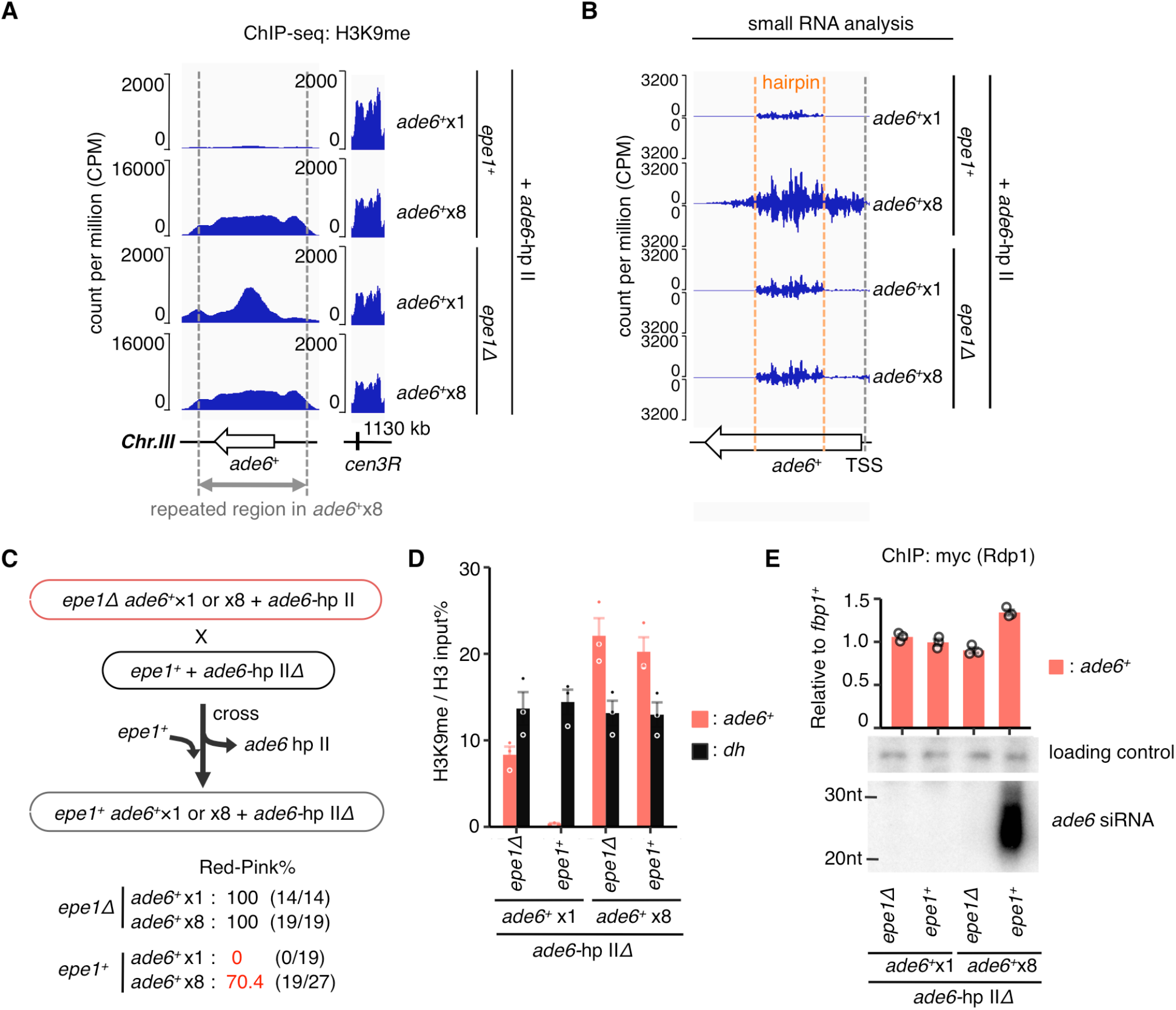
Epe1 plays opposing roles in epigenetic inheritance of H3K9me in a repeat-dependent manner. (*A*) ChIP-seq and (*B*) Small RNA-seq of *epe1Δ* cells harboring *ade6*-hp II. For comparison, results of *epe1*^*+*^ cells derived from Fig. 2F and Fig. 3A are also shown. (*C*) Red *epe1Δ* cells harboring *ade6*^*+*^x1 or *ade6*^*+*^x8 alleles that were heterochromatinized by *ade6*-hp II, were crossed with *epe1*^*+*^ *ade6*-hp II*Δ* cells to segregate *ade6*-hp II and concurrently return *epe1*^*+*^. The percentage of clones that formed red-pink (*ade6*-repressed) colonies without *ade6*-hp II was evaluated. These progenies were used for ChIP-qPCR of (*D*) H3K9me, and (*E*) an RDRC component Rdp1 (upper) and Northern blot analysis of *ade6* siRNA (lower). Error bars represent SEM; n = 3 biological replicates.

Given that the establishment of repeat-induced RNAi depends on the copy number of repeated genes as well as Epe1, these results prompted us to assume that Epe1 promotes *cis*-acting RNAi in repeat-dependent manner, and that this is why *trans*-acting RNAi failed to establish *cis*-acting RNAi at the *ade6*^*+*^x1 allele, even in the presence of Epe1. However, since Epe1 is localized to heterochromatin via the H3K9me binding protein Swi6/HP1 (Zofall and Grewal 2006), the low levels of H3K9me on the *ade6*^*+*^x1 allele in wild-type cells raise the alternative possibility that inefficient production of secondary siRNA on *ade6*^*+*^x1 is due to defective localization of Epe1. To distinguish these two possibilities, we next examined how Epe1 acts on ectopic heterochromatin already present at *ade6*^*+*^x1 or *ade6*^*+*^x8, respectively. Thus, *ade6*-hp II was removed from *epe1Δ* cells in which robust ectopic heterochromatin was already established on the *ade6*^*+*^x1 or *ade6*^*+*^x8 allele, and *epe1*^*+*^ was concurrently restored (Fig. 6C). As expected, without *epe1*^*+*^ restoration, the ectopic heterochromatins on both *ade6*^*+*^x1 and *ade6*^*+*^x8 allele were inherited even after the removal of *ade6*-hp (Fig. 6C,D). However, when *epe1*^*+*^ was restored to the *ade6*^*+*^x1 strain, all progenies formed white (*ade6*-expressing) colonies, and ectopic heterochromatin on *ade6*^*+*^ was completely removed (Fig. 6C,D). By contrast, when Epe1 was restored to the *ade6*^*+*^x8 strain, red-pink (*ade6*-repressed) progenies frequently emerged, and H3K9me levels of these progenies at *ade6*^*+*^ were not affected (Fig. 6C,D). Consistently, restored Epe1 promoted recruitment of RNAi components, such as the RDRC, and the autonomous production of *ade6* siRNAs was observed in these red-pink (*ade6*-repressed) progenies (Fig. 6E). These results indicate that Epe1 requires repeated genes to promote *cis*-acting RNAi, and when Epe1 fails to promote *cis*-acting RNAi, it erases heterochromatin. Thus, the success or failure of establishment of repeat-induced RNAi causes Epe1 to have opposing effects in epigenetic inheritance of H3K9me.

## Discussion

Constitutive heterochromatin in eukaryotic cells is enriched on repetitive DNA elements, although the molecular basis of this link is still not understood. In fission yeast, the RNAi pathway promotes constitutive heterochromatin formation at *dg*/*dh* elements, which exist at pericentromeres and other sites. RNAi-mediated heterochromatin formation specifically targets ncRNAs transcribed from *dg*/*dh* elements, but the mechanism that distinguishes *dg*/*dh* ncRNAs from other mRNAs is not clear. In this study, we have shown that an anti-silencing factor, Epe1, induces ncRNA transcription from widespread TSSs in *dg*/*dh* elements. These ncRNAs enable the assembly of the RNA machinery, thereby facilitating *cis*-acting RNAi at constitutive heterochromatin. Similar to *dg*/*dh* elements with widespread TSSs, we found that tandemly repeated euchromatic mRNA genes can also establish *cis*-acting RNAi to autonomously maintain heterochromatin there. This repeat-induced RNAi enabled us to reveal that, while Epe1 primarily functions as a heterochromatin eraser, it also promotes *cis*-acting RNAi when repeated genes underlie silent heterochromatin.

The repeat-induced RNAi described here is the first RIGS phenomenon reported in fission yeast. Previous studies of RIGS in other organisms have proposed several models to explain this universal phenomenon. In these models, tandemly repeated genes cause gene silencing directly because abnormalities derived from repetition, such as DNA-DNA interactions, aberrant transcription, or excess of a gene dosage threshold, are thought to trigger RIGS (Gladyshev and Kleckner 2017; Luo and Chen 2007; Schubert et al. 2004). By contrast, in this study, we show that the presence of tandemly repeated genes itself does not induce heterochromatin formation but instead provides an environment suitable for *cis*-acting RNAi to maintain heterochromatin autonomously. The establishment of repeat-induced RNAi depends on repeated gene copy number, as has been reported for RIGS in other organisms. Furthermore, our results showed that the effect of repeated genes results from the clustering of multiple genes in one location, and that repeat-induced RNAi requires an anti-silencing factor, Epe1. Thus, repeat-induced RNAi in fission yeast identifies a novel model of RIGS, whereby tandemly repeated genes underlying silent heterochromatin enable an anti-silencing factor to supply enough RNA templates for the assembly of the RNAi machinery, thereby facilitating *cis*-acting RNAi (Fig. 7). The requirement for repeated genes clustered in one location suggests that such a situation enables Epe1 to increase the local concentration of nascent RNA without the robust transcription that can disrupt silent heterochromatin (Shimada et al. 2016). This provides an explanation for the discrepancy between silent heterochromatin formation and the transcription necessary for the RNAi pathway. It has been reported that RNAi factors are also required for RIGS of repetitive transgenes in *D*.*melanogaster* and *C*.*elegans*, suggesting that similar mechanisms also exist in these higher eukaryotes (Pal-Bhadra et al. 2004; Kim et al. 2005).

**Figure 7.**
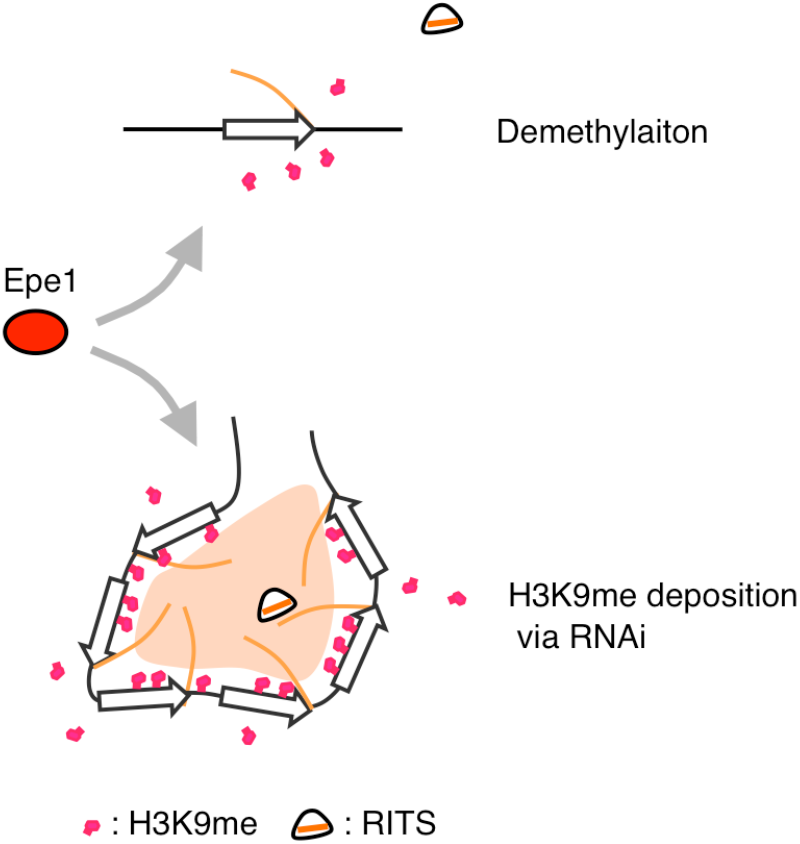
Model for repeat-induced RNAi via the anti-silencing factor Epe1. When a single target gene underlies silent heterochromatin, the number of RNA templates induced by an anti-silencing factor, Epe1, is not sufficient to assemble the RNAi machinery, thus resulting in removal of H3K9me. By contrast, repeated target genes enable Epe1 to supply enough RNA templates to assemble the RNAi machinery, leading to replenishment of H3K9me and autonomous maintenance of heterochromatin.

Notably, we found that, while Epe1 OP caused hyper-activation of the RNAi pathway at *dg*/*dh* elements, it did not cause hyper-activation of repeat-induced RNAi at *ade6*^*+*^x8 but instead removed H3K9me (Supplemental Fig. S10). This result clearly indicates that RNAi-dependent heterochromatin formation depends on the balance between replenishment and removal of H3K9me via the RNAi pathway and Epe1, respectively. Under normal levels of Epe1, the replenishment and removal of H3K9me associated with repeat-induced RNAi at *ade6*^*+*^x8 are balanced; however, when Epe1 is overexpressed, this balance is disrupted, and removal of H3K9me becomes dominant. Consistently, despite the hyper-activation of RNAi, H3K9me levels at *dg*/*dh* elements were not increased by Epe1 OP (Fig. 1A,B;Supplemental Fig. S1H). On the contrary, higher levels of Epe1 OP reduced H3K9me at *dg*/*dh* elements (Supplemental Fig. S1F) (Zofall and Grewal 2006). These results suggest that the same is true for the RNAi pathway at *dg*/*dh* elements. The number of repeat copies, the amount of Epe1, and possibly the nature of the target transcript itself will affect this balance, as discussed below.

Previous studies using *trans*-acting RNAi demonstrated that disruption of the 3’UTR region of a target gene or mutation of the highly conserved RNA polymerase-associated factor 1 complex (Paf1C), which impairs transcription termination and nascent transcript release, can promote heterochromatin formation via RNAi, even with a single target gene (Kowalik et al. 2015; Yu et al. 2014). These results suggest that retention of the nascent transcript on chromatin promotes RNAi-mediated heterochromatin formation. In particular, in the Paf1C mutants, once *trans*-acting RNAi establishes heterochromatin on a single *ade6*^*+*^ gene, it appears to be maintained during mitosis/meiosis without a supply of siRNAs in *trans* (Kowalik et al. 2015).

This result suggests that autonomous *cis*-acting RNAi is established at a single *ade6*^*+*^gene in the Paf1C mutants, analogous to the repeat-induced RNAi at *ade6*^*+*^x8 seen in this study. We speculate that perturbation of transcription termination and nascent transcript release may produce a similar effect as tandemly repeated genes, namely, an increase in the local concentration of nascent RNAs in the vicinity of chromatin. On the other hand, a recent study using *trans*-acting RNAi derived from a euchromatic fragment integrated into pericentromeric constitutive heterochromatin instead of hairpin RNA, showed that, once established, heterochromatin at the corresponding region of the chromosome arm is autonomously maintained in the absence of *trans*-acting RNAi (Yu et al. 2018). The maintenance of this ectopic heterochromatin formation is accompanied by siRNA production from adjacent genes located in corresponding regions. Although the authors did not test whether Epe1 is required to produce these multiple siRNAs, this result prompts us to speculate that, when multiple siRNAs target cognate multiple genes that exist in close proximity on the genome, these target genes behave like repeated genes and establish autonomous *cis*-acting RNAi.

Our 5’RACE analysis revealed that *dg*/*dh* elements at the mating-type locus (*cenH*) include Epe1-independent TSSs, i.e., silencing-resistant TSSs, as well as Epe1-dependent widespread TSSs. These silencing-resistant TSSs are probably counterparts of the one previously identified at the pericentromeric *dg*/*dh* element (Djupedal et al. 2005; Buscaino et al. 2013). Our truncation analysis of a *cenH* fragment with a plasmid-based minichromosome indicates that a truncated *cenH* fragment that contains only widespread TSSs can establish heterochromatin. This result suggests that silencing-resistant TSSs are dispensable for RNAi-mediated heterochromatin formation. On the other hand, the *ade6*^*+*^ gene, which was used as a reporter gene for the repeat-induced RNAi system in this study, is a euchromatic mRNA gene without introns, and no obvious antisense transcripts specific for the *ade6*^*+*^ repeat were detected. Therefore, the establishment of repeat-induced RNAi at *ade6*^*+*^x8 suggests that previously reported properties of *dg*/*dh* ncRNAs, such as silencing-resistant TSSs (Djupedal et al. 2005; Buscaino et al. 2013), bidirectional transcription (Volpe et al. 2002), and splicing (Bayne et al. 2008), are dispensable for heterochromatin formation via RNAi. These properties may instead be required for production of primary siRNAs, which were replaced by hairpin RNAs in this study, or they may enhance the efficiency of assembly of the RNAi machinery at *dg*/*dh* ncRNAs. Indeed, previous studies have shown that *dg*/*dh* elements produce primary siRNAs in the absence of the RDRC (Yu et al. 2014). Furthermore, although widespread TSSs were detected throughout *dg*/*dh* elements in our study, *dg*/*dh* siRNAs are only produced from a limited portion of *dg*/*dh* elements, so-called siRNA hot spots (Djupedal et al. 2009). These facts suggest that *dg*/*dh* elements have additional mechanisms that produce primary siRNAs and/or promote the assembly of the RNAi machinery on a subpopulation of *dg*/*dh* ncRNAs.

Constitutive heterochromatin is enriched not only on repetitive transgenes but also on repeated DNA sequences, such as a satellite repeat at a pericentromere. It is not clear whether the mechanism responsible for RIGS is also applicable to repeated DNA sequences, which do not encode mRNA genes. In this study, we show that RNAi-mediated heterochromatin formation at pericentromeric *dg*/*dh* elements is also dependent on Epe1, and that the *dg*/*dh* elements encode widespread TSSs with a preference for a core promoter structure identical to that of mRNAs. These results suggest that *dg/dh* elements also function as a variant of repeated mRNA genes; ncRNAs transcribed from closely placed TSSs will behave like tandemly repeated mRNA genes because such transcripts have the same sequences as each other. Thus, the flexibility of repeat-induced RNAi indicated by *dh*/*dh* elements prompts us to speculate that a simple repeated DNA sequence that does not encode mRNAs could also produce the same effect, as long as it is transcribed. Although no Epe1 orthologs have thus far been identified in higher eukaryotes, anti-silencing mechanisms that direct transcription from silent heterochromatin have been reported (Andersen et al. 2017; Law et al. 2013). The dual and opposing roles of an anti-silencing factor discovered in this study provide a novel model to explain why repetitive DNA elements are linked to constitutive heterochromatin in eukaryotic cells.

## Materials and methods

### *Schizosaccharomyces pombe* strains and genetic manipulations

All strains used in this study are described in Supplemental Table S1. A PCR-based method was used for deletion or epitope tagging of target genes. All integrations were confirmed by PCR or Southern blot. Silencing assays for *ade6*^*+*^ gene were performed with PMG-based synthetic media containing a limited concentration of adenine (10 mg/L), as the indicator plate. To evaluate the efficiency of establishment or maintenance of the *ade6*-repressed state, red-pink (*ade6*-repressed) colony formation from white (*ade6*-expressing) or red (*ade6*-repressed) originator were assessed, respectively. For Epe1 OP, the endogenous *epe1*^*+*^ promoter was replaced with Purg1 (Watt et al. 2008). For comparison between Epe1 OP and higher levels of Epe1 OP (expression from multi-copy plasmid pREP41) (Maundrell 1993), see Supplemental Fig. S1D-G.

### RT-PCR and qRT-PCR

Total RNA was extracted using the hot phenol method and treated with recombinant DNase I (Takara, cat# 2270A) in the presence of RNasin Plus (Promega, cat# N2611).

PrimeScript Reverse Transcriptase (Takara, cat# 2680A) was used for reverse transcription according to the manufacturer’s instructions. The primers used for analysis are listed in Supplemental Table S2.

### Northern blot analysis of siRNA

Small RNAs were extracted using the mirVana miRNA Isolation Kit (Ambion, cat# AM1561) according to the manufacturer’s instructions. Small RNAs were separated on a 15% sequencing gel and transferred to Amersham Hybond-N+ membranes (GE Healthcare, cat# RPN303B) using a Trans-blot SD semi-dry electrophoretic transfer cell (Bio-Rad, cat# 170-3940). After UV crosslinking, probes labeled with [α-^32^P] dCTP by random priming (Takara, cat# 6045) were hybridized to the membrane in PerfectHyb Plus Hybridization buffer (Sigma, cat# H7033) at 42°C overnight. After washing with 2× SSC 0.1% SDS buffer at 42°C, the membrane was exposed to an imaging plate for 1–2 days. An oligonucleotide probe against snoRNA58 labeled with [γ-^32^P] ATP by T4 polynucleotide kinase (Takara, cat# 2021A) was used as a loading control. The primers used for this analysis are listed in Supplemental Table S2.

### Small RNA-seq

After electrophoresis of small RNAs on a 15% sequencing gel, gel fractions corresponding to 20–30 nt were harvested and crushed in extraction buffer (0.3M NaCl). After overnight rotation at 4°C, the small RNAs were recovered by ethanol precipitation using Ethachinmate (NIPPON GENE, cat# 318-01793) as a carrier. The small RNA (smRNA) libraries were constructed using the SMARTer smRNA-seq kit for Illumina (CLN cat#635029) according to the manufacturer’s instructions. After purification by AMPure XP (Beckman Coulter, cat# A63882), smRNA libraries were sequenced on the Illumina HiSeq 4000 system (single-end, 51 bp) or HiSeqX system (paired-end, 151 bp). Reads (Read 1 in the case of paired-end) were first trimmed for adaptors and A-tailing incorporated during library construction using the cutadapt program (Martin, 2011), with the following parameters; -m 15 -u 3 -a AAAAAAAAAA. Using bowtie (version 1.2.1.1) with -M 1 --best parameters, the trimmed reads were mapped on the modified *S. pombe* genome, in which the native *ade6*^*+*^ ORF is deleted, and the *ura4*^*+*^ gene is replaced by the *ade6*^*+*^x1 construct. The Bedgraph files for visualizing in the IGV were produced with the “genomecov” function of the BEDTools program (version 2.26), which were then normalized by 1 million total mapped reads.

### ChIP-qPCR and ChIP-seq

ChIP experiments were performed as described previously (Kawakami et al. 2012). The antibodies used were H3K9me2 antibody (mAbProtein, m5.1.1; a gift from Takeshi Urano, Shimane University), anti-Histone H3 antibody (Millipore, cat# 07-690), ANTI-FLAG M2 antibody (Sigma, cat# F1804), the anti-Myc tag antibody clone 4A6 (Millipore, cat# 05-724), and the anti-RNA polymerase II antibody clone CTD4H8 (Millipore, cat# 05-623). Note that H3K9me2 antibody (mAbProtein, m5.1.1) recognizes H3K9 mono-, di-, and trimethylation (Takeshi Urano, personal communication). Enrichment relative to the euchromatic *fbp1*^*+*^ was evaluated for ChIP of RNAi factors, CLRC component, and Pol2. For ChIP of H3K9me, the amount of immunoprecipitated H3K9me relative to the input whole cell extract (input%) was normalized to that of H3 because of its low background signals at *fbp1*^*+*^. In Fig. 1 and Supplemental Fig. S1, the amount of immunoprecipitated H3K9me relative to the input whole cell extract (input%) was normalized to that of the wild type. The primers used for analysis are listed in Supplemental Table S2. For ChIP-seq, the ChIP libraries were prepared with a KAPA Hyper Prep Kit (KAPABIOSYSTEMS, cat#KK8504), according to the manufacturer’s instructions. The libraries were sequenced on the Illumina HiSeq 2500 system (single-end, 50 bp). The sequenced reads were mapped on the modified *S. pombe* genome using bowtie (version 1.2.1.1). The data were processed by SAMtools (version 1.9) and IGVTools to make tdf files that were visualized in the IGV.

### CAGE-seq

CAGE library preparation, sequencing, mapping, and gene expression analysis were performed by DNAFORM (Yokohama, Kanagawa, Japan). First-strand cDNAs were transcribed to the 5’-end of capped RNAs and attached to CAGE “bar code” tags, and the sequenced CAGE tags were mapped to the *S. pombe* ASM294v2 genome using BWA software (0.7.15-r1140), after discarding ribosomal or non-A/C/G/T base-containing RNAs. CAGE-tag data without clustering were used for visualization of TSSs and gene expression profiles to avoid the loss of signals derived from widespread TSSs at *dg*/*dh* elements. To confirm reproducibility, CAGE-seq was repeated twice for total RNA from Epe1 OP cells, and representative data are presented. For additional analyses of CAGE-seq with Weblogo3 (Crooks et al. 2004), see also Supplemental methods.

## Data availability

CAGE-seq, ChIP-seq, and smRNA-seq data have been submitted to the DDBJ (https://www.ddbj.nig.ac.jp) under accession nos. DRA006868, DRA013983, DRA013984, DRA014003, DRA014324.

## Acknowledgments

We thank R. Allshire for providing strains and plasmids; H. Kato for technical advice; T. Urano for providing antibodies; DNAFORM for CAGE-seq; H. Masumoto, M. Siomi, S. Yamanaka, and J. Nakayama for critical reading; our laboratory members for helpful discussions; T. Matsumoto and his lab members for mentoring of TA; and A. Kanji, Keiko, and Yoko for their support of TA. This work was supported by funding to YM from JSPS Grant-in-Aid for Scientific Research (A) 10159209, 12206045, and MEXT Grant-in-Aid for Scientific Research on Priority Areas 21247001, and to SI from MEXT Grant-in-Aid for Transformative Research Areas (A) JP20H05913 (SI), and to TA from JSPS (DC2).

## Author contributions

Conceptualization: TA, YM

Investigation: TA, SI, TK, HA

Writing – original draft: TA, YM

Writing – review & editing: TA, YM

