## Supplemental Figs for "Tandemly repeated genes promote RNAi-mediated heterochromatin formation via an anti-silencing factor Epe1 in fission yeast"

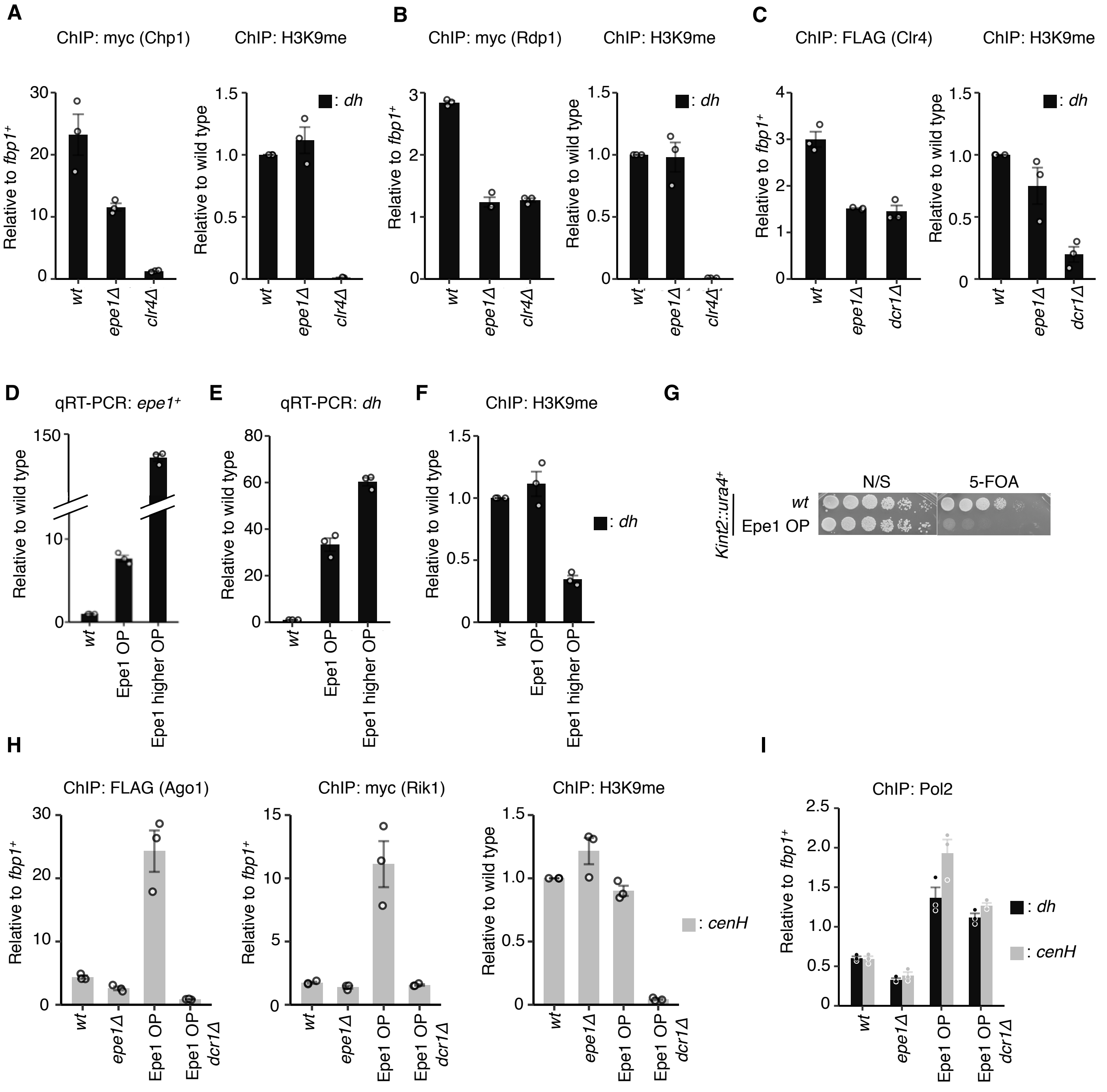


**Supplemental Figure S1. Epe1 promotes the assembly of the RNAi machinery on constitutive heterochromatin.** (*A*) ChIP-qPCR of the RITS complex component, Chp1; (*B*) the RDRC component, Rdp1; and (*C*) the CLRC component, Clr4, at pericentromeric *dh*. Because each *epe1Δ* clone shows different levels of H3K9me (Trewick et al. 2007), results of simultaneously performed ChIP-qPCR of H3K9me are shown side by side. (*D*)(*E*)(*F*), comparison between Epe1 OP (endogenous *epe1*$^+$ promoter, replaced by Purg1 (Watt et al.

2008)) and higher levels of Epe1 OP (expression from multi-copy plasmid pREP41(Maundrell 1993)). (*D*) qRT-PCR of *epe1*$^{+}$ transcripts relative to wild type. (*E*) qRT-PCR of *dh* transcripts relative to wild type. (*F*) ChIP-qPCR of H3K9me at *dh*. (*G*) Silencing assays of Epe1 OP strains, in which a reporter gene, *ura4*$^{+}$, is inserted within heterochromatin at the mating-type locus (*Kint2::ura4*$^{+}$) (see also Supplemental Fig. S3A). To assess *ura4*$^{+}$ gene silencing, 4.5-fold serial dilutions of log-phase cells were spotted onto each medium based on Edinburgh minimal medium (EMM). FOA; the EMM 5-FOA plate contains 0.2% 5-fluoroorotic acid (FOA), which is toxic to cells expressing *ura4*$^{+}$. N/S, non-selective. (*H*) Epe1 OP promotes the assembly of RNAi machinery on *dg*/*dh* elements at the mating-type locus (*cenH*). ChIP-qPCR analyses of Ago1, the CLRC component, Rik1, and H3K9me at *cenH* are shown. (*I*) ChIP-qPCR of Pol2 at *dh* or *cenH* elements. Error bars represent SEM; n = 3 biological replicates.

Asanuma_Fig_S2

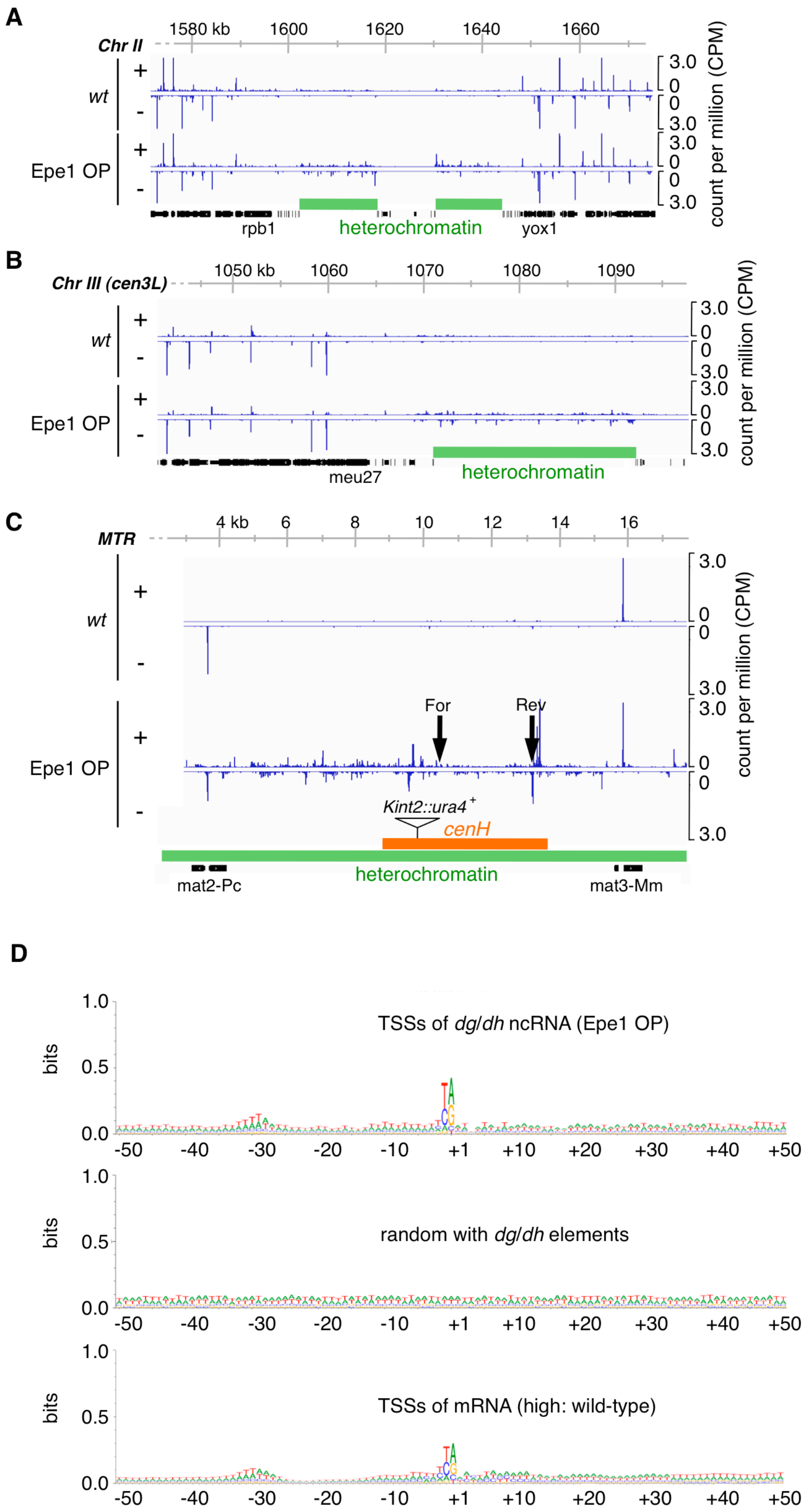

**Supplemental Figure S2. Epe1 overproduction promotes ncRNA transcription from TSSs that are widespread across constitutive heterochromatin.** (*A*)(*B*)(*C*) CAGE reads of the indicated strains at (*A*) centromere II, (*B*) centromere III, and (*C*) the mating-type locus (MTR) are shown. Green horizontal bars indicate heterochromatin region. An orange horizontal bar indicates the *cenH* region analyzed in Supplemental Fig. S3. Arrowheads indicate the position of forward (For) and reverse (Rev) major transcription start sites (TSSs) identified by 5'RACE analysis (see also Supplemental Fig. S3). (*D*) Consensus sequence analysis of widespread TSSs in *dg*/*dh* elements. The sequences of the ±50 nt flanking TSSs in *dg*/*dh* elements, which showed more than 0.05 CPM under Epe1 OP, were analyzed using WebLogo (Crooks et al. 2004). The sequences of ±50 nt randomly picked from *dg*/*dh* elements were also analyzed as a control. TSSs of mRNA in wild-type cells were categorized into three groups according to expression strength. The sequences of ±50 nt flanking unique TSSs of mRNAs categorized as having high expression were analyzed. The horizontal axis indicates relative position with respect to TSS (+1). TSSs of *dg*/*dh* elements and euchromatic genes commonly exhibit a preference for the consensus initiator (Y/R) sequence at -1/+1 positions and an A/T rich region 25-32 nt upstream.

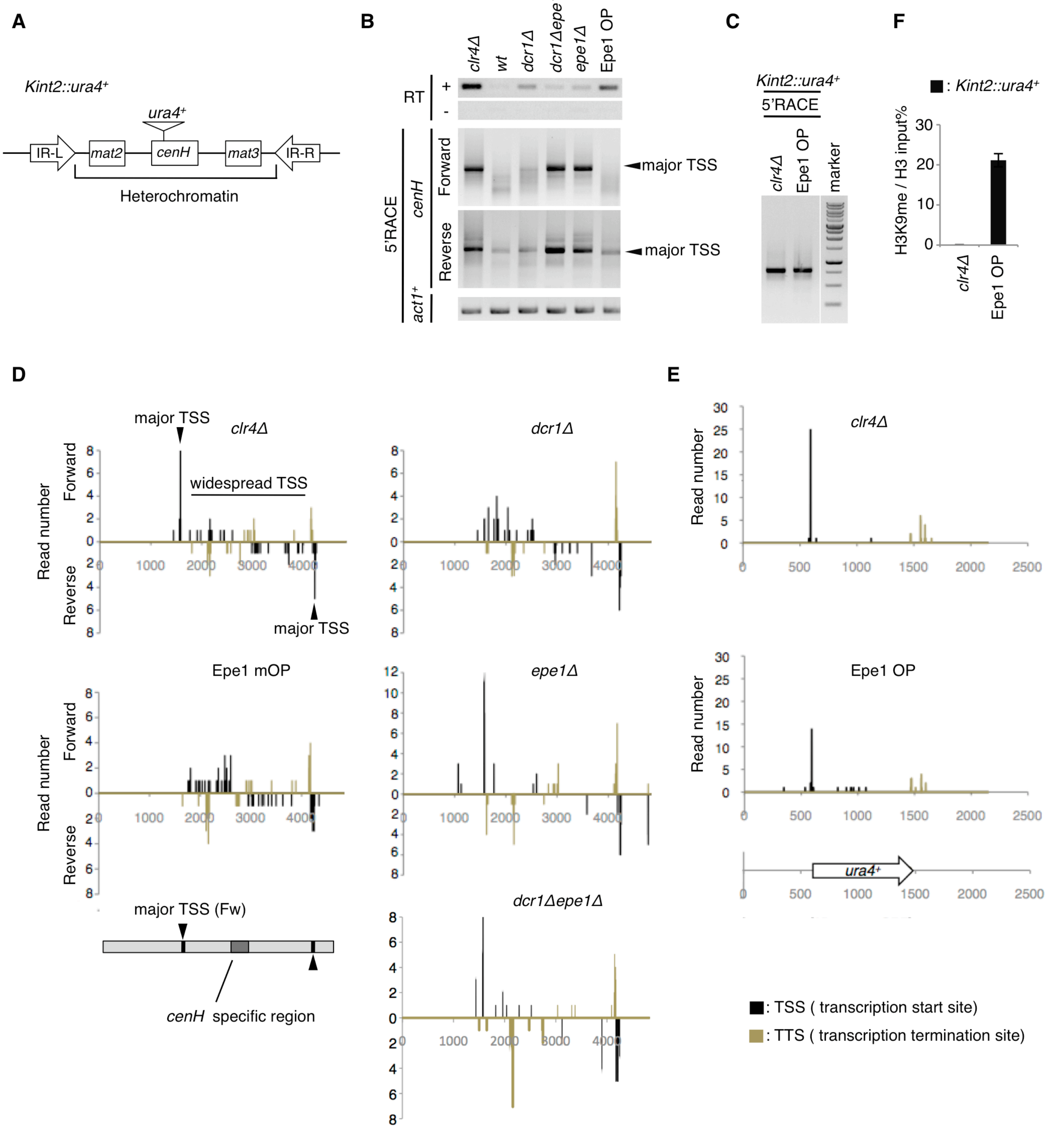


**Supplemental Figure S3. RACE analysis of *dg*/*dh* elements at the mating-type locus (*cenH*) and inserted reporter gene, *ura4*$^{+}$.** (*A*) Schematic diagram of the mating-type locus containing *cenH* and inserted *ura4*$^{+}$ (*Kint2::ura4*$^{+}$) (Jia et al. 2004). The *cenH* locus has a specific insertion that enables *cenH* ncRNAs to be distinguished from other *dg*/*dh* ncRNAs. (*B*)(*C*) Representative image of 5'RACE for (*B*) forward and reverse *cenH* ncRNAs, *act1*$^{+}$ mRNAs, and (*C*) *Kint2::ura4*$^{+}$. RT-PCR of *cenH* ncRNAs using oligodT primers is also shown at the top of (*B*). (*D*)(*E*) TSSs identified by 5'RACE were mapped to genomic sequences of (*D*) *cenH* or (*E*) *ura4*$^{+}$. Note that TSSs could not be identified with wild-type cells, because their RACE products were poorly cloned. There are two types of TSS for *cenH* ncRNAs: major TSSs and widespread TSSs (*D*, *clr4Δ*). Notably, while *epe1Δ* inactivated widespread TSSs, Epe1 OP caused their hyper-activation, resulting in smears on the gel (*B*). By contrast, at *Kint2::ura4*$^{+}$, Epe1 OP did not cause smearing but activated TSSs corresponding to those observed in the absence of heterochromatin (*clr4Δ*). Note that Epe1 OP induces the expression of *Kint2::ura4*$^{+}$ in the presence of heterochromatin (*F*). Transcription termination sites (TTSs) identified by 3'RACE are also plotted in (*D*) (*E*).

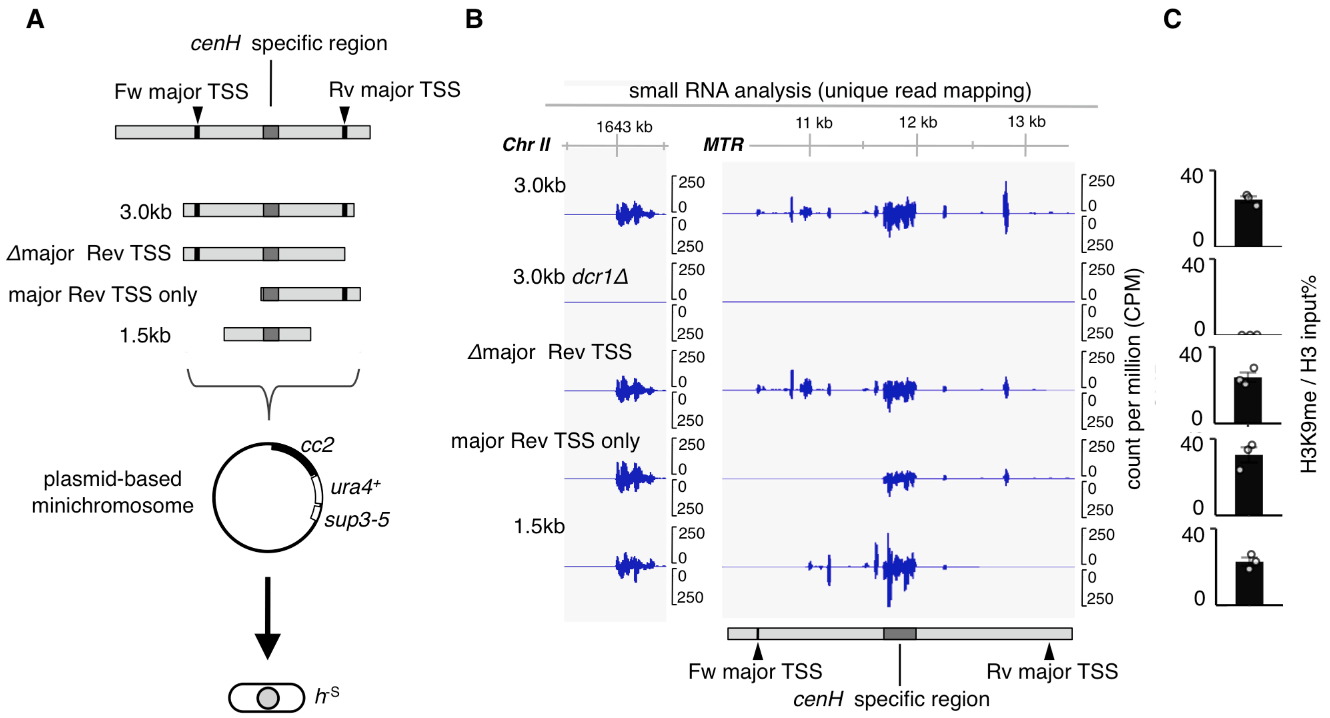


**Supplemental Figure S4. A *cenH* fragment that contains only widespread TSSs can establish heterochromatin.** (*A*) Diagram of the experimental scheme for truncation analysis of the *cenH* fragment. The positions of forward and reverse major TSSs identified by 5'RACE (see also Supplemental Fig. S3) are indicated. The position of the *cenH*-specific region, which is utilized to evaluate siRNA production and H3K9me accumulation, is also marked. To determine which TSS elements are required for the RNAi, a series of truncated *cenH* fragments were cloned into plasmid-based minichromosomes (Buscaino et al. 2013) and transformed into the $h^{-S}$ strain, in which the native *cenH* region at the mating-type locus is completely lost. This enabled evaluation of siRNA production and H3K9me accumulation at truncated *cenH* fragment on minichromosomes. (*B*) Small RNA-seq mapping to part of the chromosome II pericentromere and *cenH* fragment is shown. To evaluate production of siRNAs on truncated *cenH* fragments easily, only uniquely mapped reads were extracted. Such extraction omits reads that are indistinguishable from those derived from redundant *dg*/*dh* elements at the pericentromere. (*C*) ChIP-qPCR of H3K9me using *cenH*-specific region primers. Error bars represent SEM; n = 3 biological replicates. These results indicate that a 1.5 kb fragment containing only widespread TSSs can establish heterochromatin.

Asanuma_Fig_S5

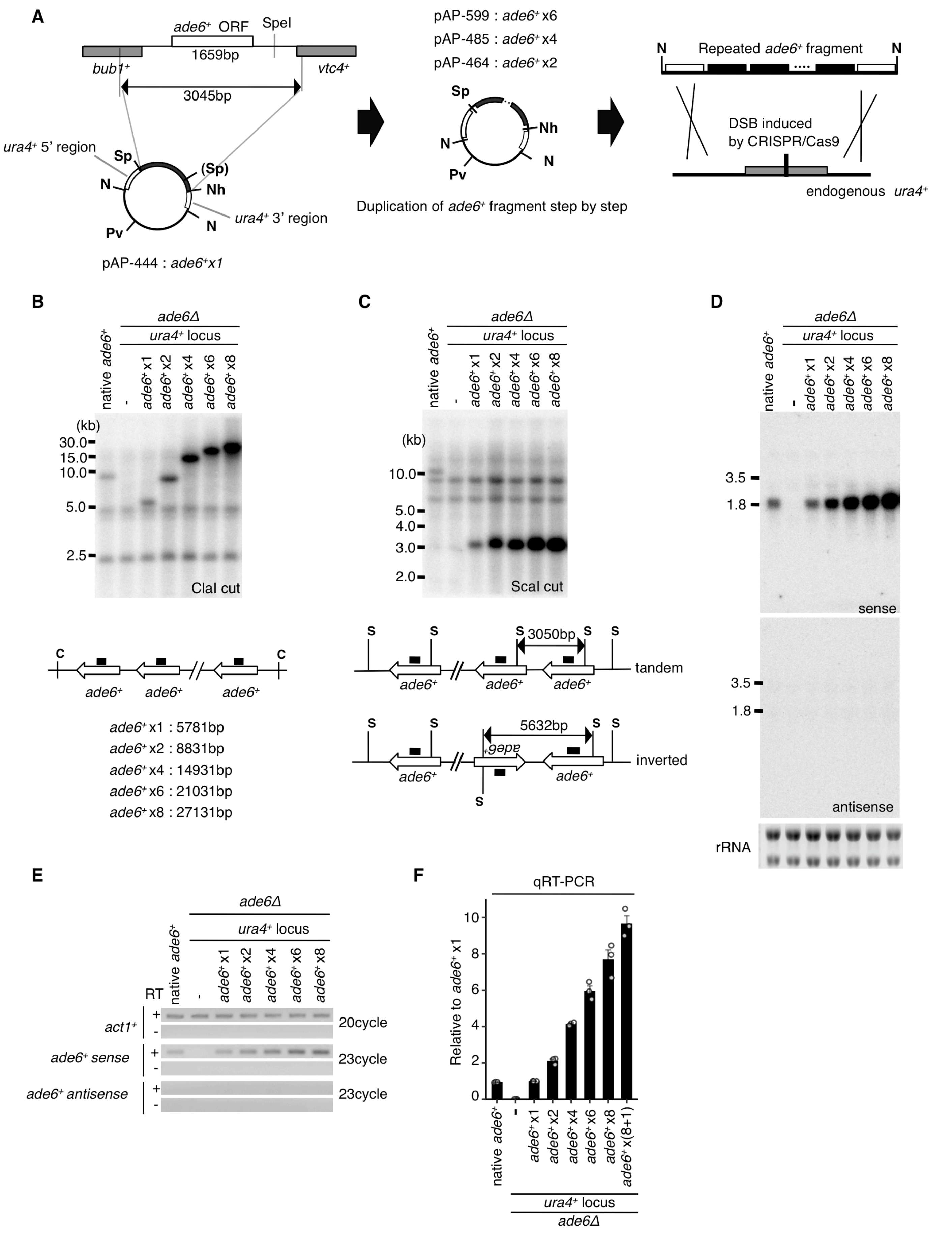

**Supplemental Figure S5. Generation of *ade6*$^+$ repeat strains.** (*A*) Schematic diagram illustrating the construction of *ade6*$^+$ repeat strains. The *ade6*$^+$ fragment was cloned between *ura4*$^+$ homologous sequences on a plasmid and then duplicated stepwise (see also Supplemental methods). A CRISPR/Cas9-dependent double strand break (DSB) at the endogenous *ura4*$^+$ locus was employed to promote integration of the *ade6*$^+$ repeat fragments by homologous recombination. The restriction sites used for construction and recovery of *ade6*$^+$ repeat fragments are shown: N, NotI; Sp, SpeI; Nh, NheI; and Pv, PvuI. Parentheses indicate the restriction site destructed by point mutation. (*B*) (*C*) Southern blot analyses of strains harboring *ade6*$^+$ repeats at the *ura4*$^+$ locus using an *ade6*$^+$ probe (black rectangle). Note that endogenous *ade6*$^+$ was deleted in these strains. Genomic DNA was digested with (*B*) ClaI or (*C*) ScaI to evaluate the length of the *ade6*$^+$ repeat and its arrangement, respectively. The diagrams below the blots illustrate the scheme for each Southern blot. (*D*) Northern blot analysis of strains harboring *ade6*$^+$ repeats at the *ura4*$^+$ locus using sense and antisense probes for *ade6*$^+$. Wild type and *ade6Δ* strain were used as positive and negative controls, respectively. rRNA stained with ethidium bromide is shown as a loading control. (*E*) RT-PCR analysis of *ade6*$^+$ transcripts using strand-specific primers for reverse transcription in the presence (+) or absence (-) of reverse transcriptase (RT). No antisense transcripts specific for the *ade6*$^+$ repeat were detected even if the number of PCR cycles was increased (data not shown). (*F*) qRT-PCR for *ade6*$^+$ mRNA using oligodT primers for reverse transcription. Error bars represent SEM; n = 3 biological replicates.

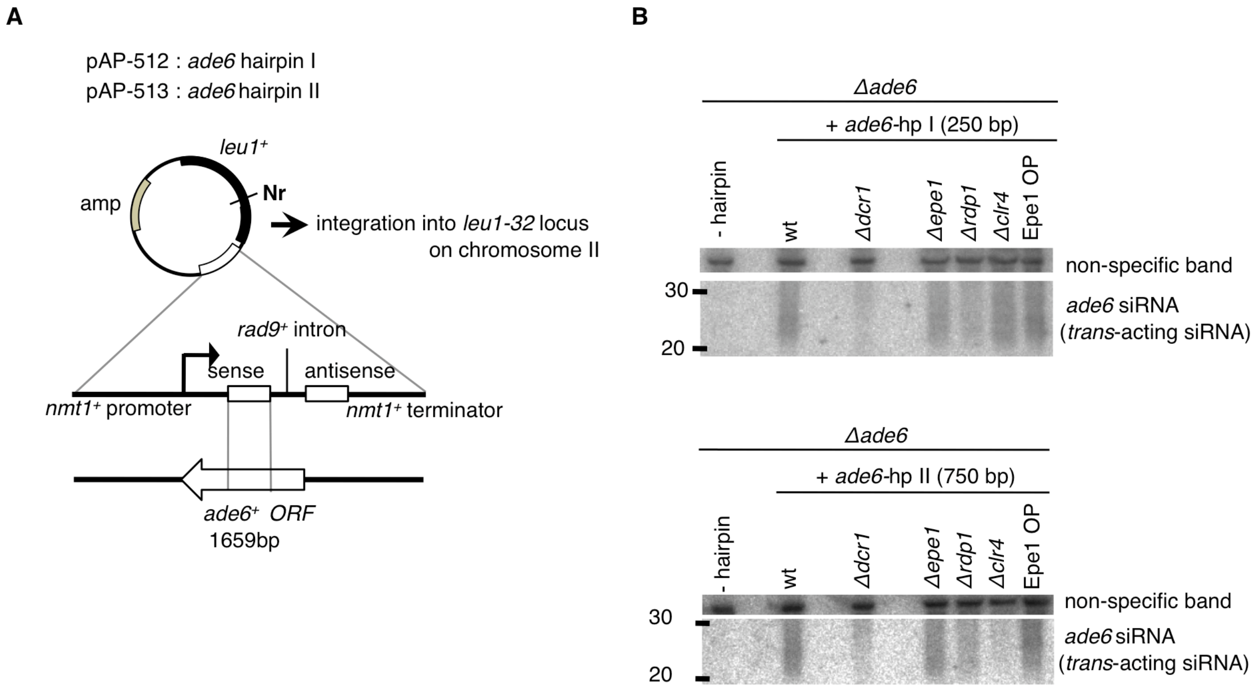


**Supplemental Figure S6. Design of *ade6* hairpin RNA constructs that drive *trans*-acting RNAi.** (*A*) Schematic diagram of *ade6* hairpin RNA constructs. The *ade6* hairpin I (*ade6*-hp I) or *ade6* hairpin II (*ade6*-hp II) construct includes a 250 bp or 750 bp fragment corresponding to 621-871 nt or 386-1132 nt of the *ade6*$^{+}$ ORF, respectively. A spacer sequence derived from the first intron of *rad9*$^{+}$ separates the inverted *ade6*$^{+}$ fragment to form the hairpin construct. The inducible *nmt1*$^{+}$ promoter drives the expression of hairpin RNAs. The constructed plasmids are integrated into the *leu1-32* locus on chromosome II of host cells. The restriction site used for linearization is shown: Nr, NruI. (*B*) Northern blot analysis using an *ade6*$^{+}$ probe to detect *trans*-acting *ade6* siRNAs derived from *ade6*-hp I or II. A non-specific band was used as a loading control. Because these strains do not have both endogenous *ade6*$^{+}$ and the *ade6*$^{+}$ repeat allele at the *ura4*$^{+}$ locus, blots represent siRNAs derived only from hairpin RNA. Note that loss of Epe1 or Epe1 OP does not affect siRNA production from hairpin RNAs. *dcr1Δ* cells were used as a negative control.

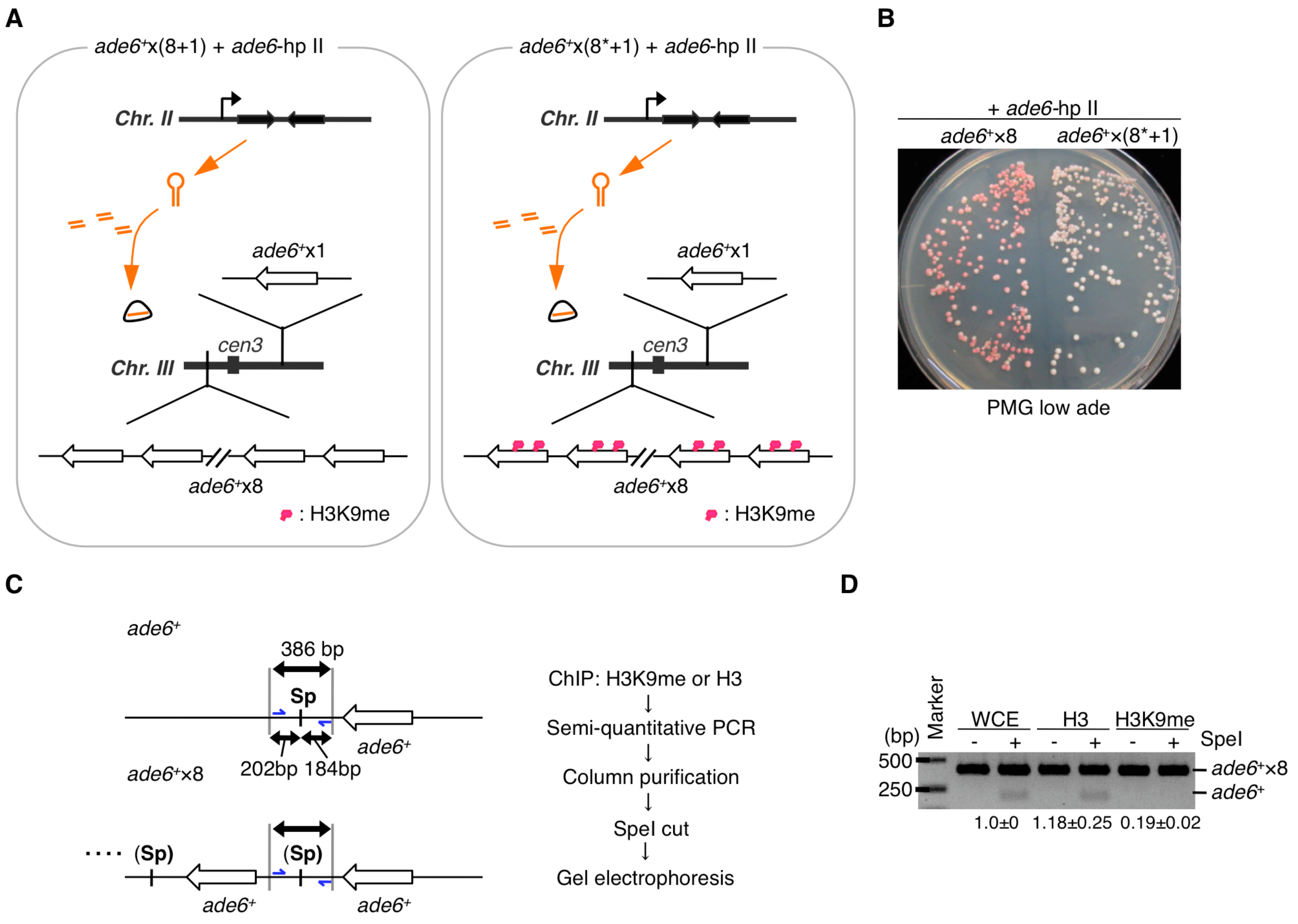


**Supplemental Figure. S7. Repeated genes do not promote RNAi-mediated heterochromatin formation by their gene dosage.** (*A*) Schematic diagram of *ade6*$^{+}$x(8+1) and *ade6*$^{+}$x(8*+1) strains in which the endogenous *ade6*$^{+}$ gene was combined with the *ade6*$^{+}$x8 allele. In *ade6*$^{+}$x(8*+1), *ade6*$^{+}$x8 allele had already been heterochromatinized by *trans*-acting RNAi (asterisk). (*B*) Silencing assay of *ade6*$^{+}$x8* and *ade6*$^{+}$x(8*+1) strains. Phenotypes after red (*ade6*-repressed) clones that were sequentially selected twice are shown. Compared with the *ade6*$^{+}$x8* strain, the red-pink phenotype of the *ade6*$^{+}$x(8*+1) strain was hardly maintained. (*C*) Diagram of the experimental scheme for semi-quantitative ChIP-PCR of H3K9me to distinguish an isolated *ade6*$^{+}$ from the *ade6*$^{+}$x8 allele. The SpeI site situated downstream of the *ade6*$^{+}$ ORF was removed by point mutation during the *ade6*$^{+}$x8 construction (see also Supplemental Fig. S5). Therefore, semi-quantitative ChIP-PCR with primers that amplify this restriction site region (blue arrows), followed by SpeI digestion, enabled immunoprecipitated DNAs of an isolated *ade6*$^{+}$ gene to be distinguished from those of the *ade6*$^{+}$x8 allele. (*D*) Semi-quantitative ChIP-PCR using the indicated antibodies was performed, and products were separated by gel electrophoresis with (+)/without (-) SpeI digestion. After electrophoresis, gels were stained with ethidium bromide, and the density of each band was measured using imageJ software. The intensity of the band derived from an isolated *ade6*$^{+}$ relative to that from the *ade6*$^{+}$x8 allele, and normalized to WCE intensity, was evaluated. Average signal intensities calculated from three independent experiments are shown. Representative image of electrophoresis is shown.

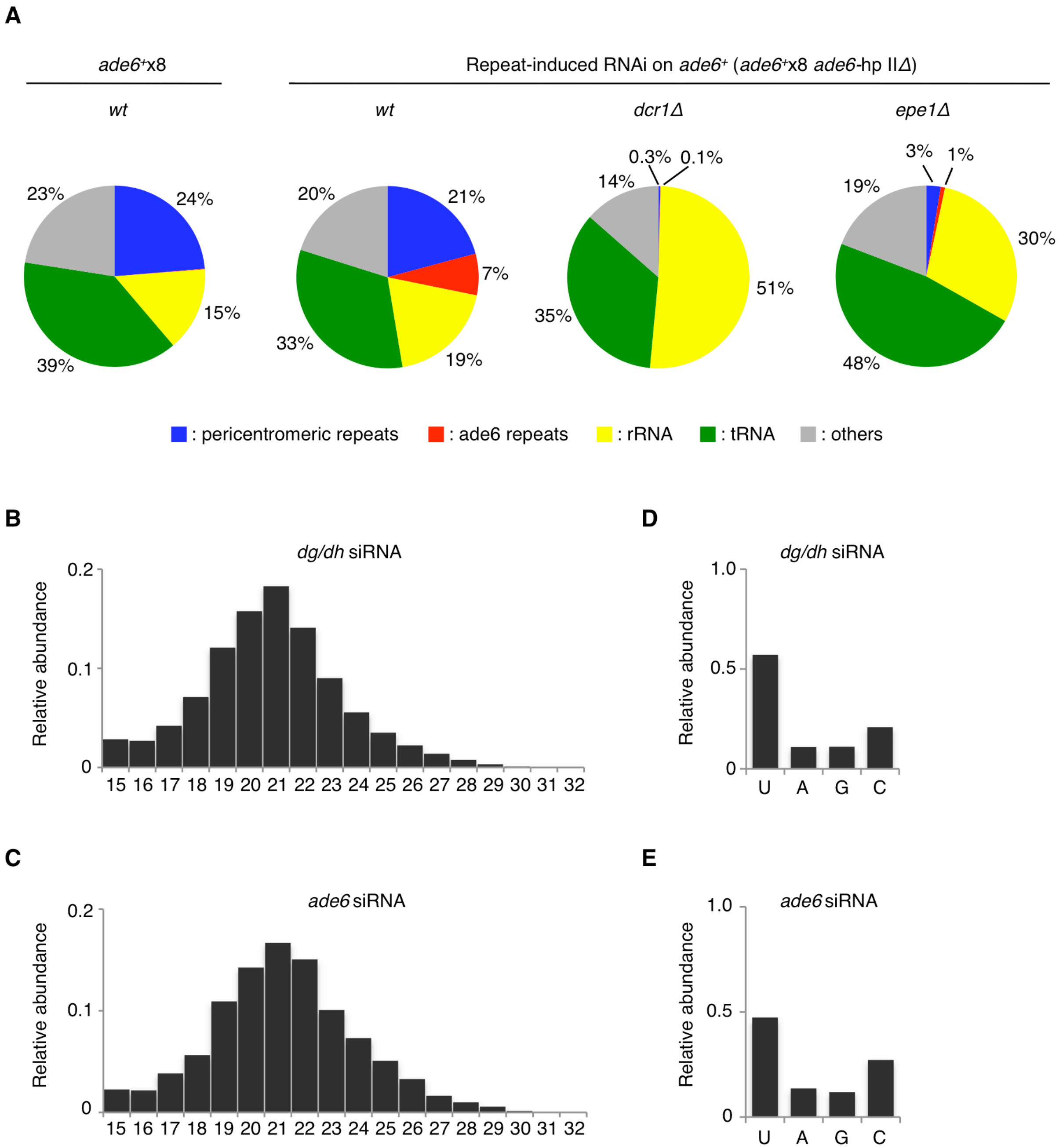


**Supplemental Figure S8. Repeat-induced RNAi on *ade6*$^{+}$x8 produces siRNAs that exhibit the same properties as native siRNAs from *dg/dh* elements.** (*A*) High-throughput sequencing was used to analyze small RNAs in the *ade6*$^{+}$x8 strain either before establishment of repeat-induced RNAi or after its establishment in combination with *dcr1Δ* or *epe1Δ*. Note that these strains do not have hairpin RNA constructs expressing *trans*-acting siRNAs. Pie charts illustrate percentages of each small RNA category relative to total small RNA reads. (*B*)(*C*)(*D*)(*E*) Small RNA reads of *ade6*$^{+}$x8 strain after repeat-induced RNAi was established were used for analysis of size distribution and 5’ nucleotide preference. (*B*)(*C*) Size distributions of small RNAs derived from (*B*) *dg*/*dh* elements or (*C*) *ade6*$^{+}$x8 are shown. Horizontal axes indicate lengths of small RNAs, and vertical axes represent relative abundance. (*D*)(*E*) Relative abundance of the first nucleotide of small RNAs derived from (*D*) *dg*/*dh* elements or (*E*) *ade6*$^{+}$x8.

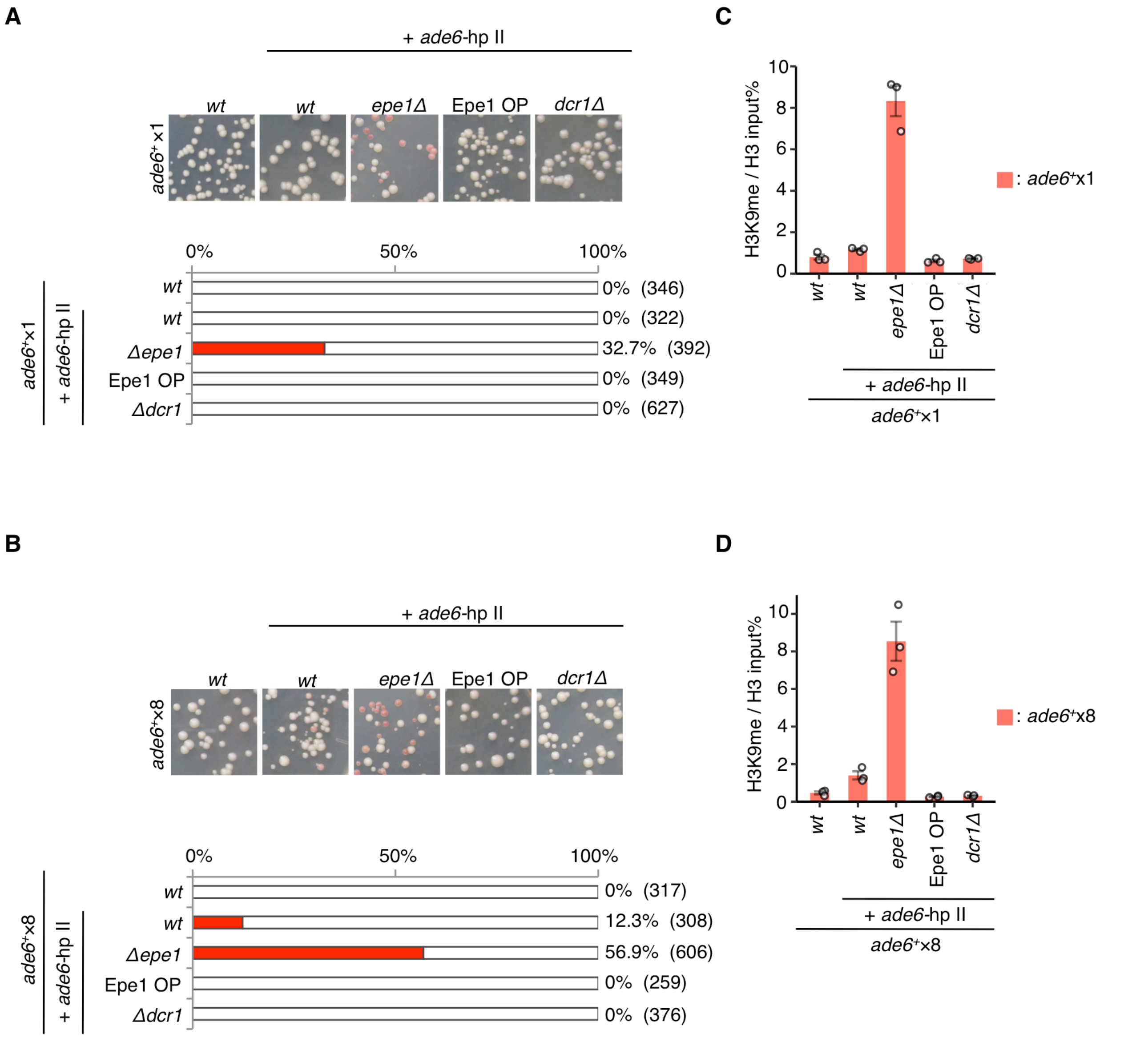


**Supplemental Figure S9. Epe1 primarily suppresses siRNA-directed heterochromatin formation.** (*A*)(*B*) Silencing assays of (*A*) *ade6*$^{+}$x1 and (*B*) *ade6*$^{+}$x8 strains harboring *ade6*-hp II. Loss of *epe1*$^{+}$ causes self-propagation of heterochromatin, which is accompanied by the appearance of the red-white variegated phenotype (see also Fig. 4E). Because such effects can perturb the evaluation of heterochromatin maintenance and establishment by the silencing assay, cells were used immediately after induction of hairpin RNA expression. Percentages of red-pink (*ade6*-repressed) colonies observed are shown in the bar graph. The numbers of colonies scored are noted in parentheses. (*C*)(*D*) ChIP-qPCR of H3K9me with the cells from (*A*) and (*B*). Error bars represent SEM; n = 3 biological replicates. Note that *epe1Δ* or Epe1 OP does not affect the amount of *trans*-acting siRNAs (Supplemental Fig. S6).

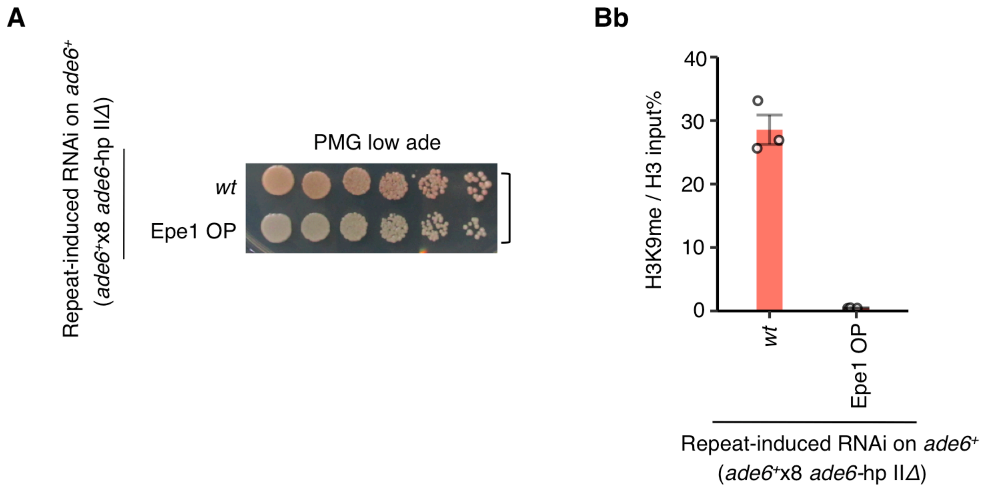


**Supplemental Figure S10. Epe1 OP removes H3K9me at *ade6*$^{+}$x8, where repeat-induced RNAi maintains heterochromatin.** (*A*) Silencing assay of the heterochromatinized *ade6*$^{+}$x8 allele combined with Epe1 OP. Red *ade6*$^{+}$x8 cells, in which repeat-induced RNAi maintains heterochromatin on *ade6*$^{+}$, were crossed with Epe1 OP cells. Wild-type cells derived from the same ascus, which inherit the cognate *ade6*$^{+}$x8 allele, are also shown as controls and indicated with close square brackets. When the heterochromatinized *ade6*$^{+}$x8 allele was combined with Epe1 OP, cells formed only white (*ade6*-expressing) colonies. (*B*) ChIP-qPCR of H3K9me shows H3K9me was completely removed from the *ade6*$^{+}$x8 allele in the Epe1 OP strain. This experiment was performed alongside those in Fig. 4F; therefore, the same data for wild-type cells (*wt*) are shown. Error bars represent SEM; n = 3 biological replicates.

Asanuma_Fig_S11

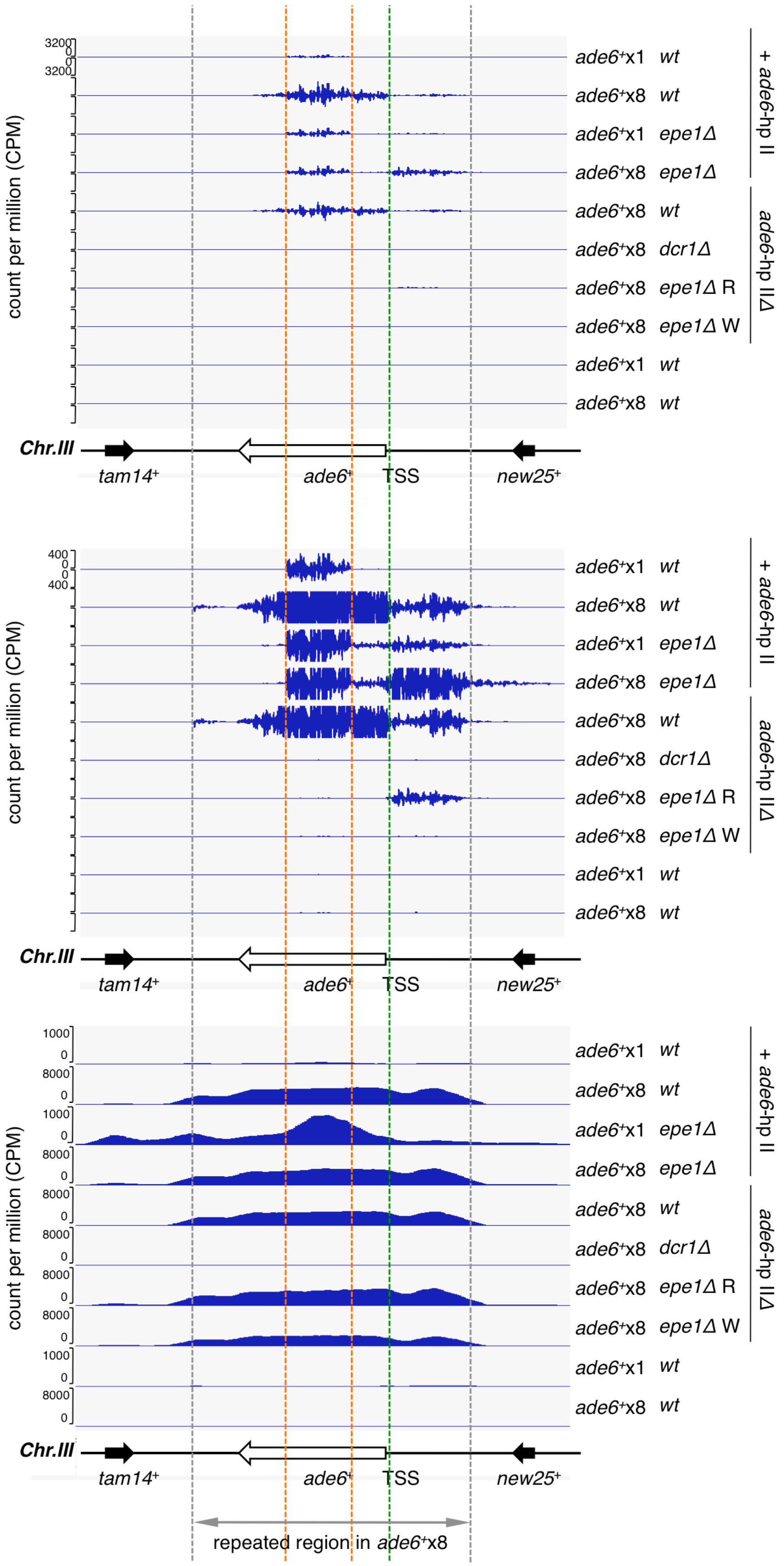

**Supplemental Fig. S11. Small RNA production in the vicinity of ectopic heterochromatin on *ade6*$^{+}$.** Small RNA-seq reads mapped in the vicinity of the *ade6*$^{+}$ gene are shown (upper). An enlarged view is shown in the middle panel. Notations of count per million are omitted for all tracks except the top one. For comparison, the results of ChIP-seq analysis for H3K9me with the same strains are shown in the lower panel. Reads from both *ade6*$^{+}$x1 or *ade6*$^{+}$x8 cells were mapped on the *ade6*$^{+}$x1 construct to facilitate visualization. Since *ade6*$^{+}$x8 strains have eight copies of the *ade6*$^{+}$ gene, ChIP-seq reads of these strains at *ade6*$^{+}$ are scaled by a factor of eight. Orange dashed lines indicate the region targeted by *ade6*-hp II (*trans*-acting RNAi). The green dashed line indicates the TSS of *ade6*$^{+}$, and gray dashed lines indicate either end of the *ade6*$^{+}$ repeat fragment. The locations and direction of *ade6*$^{+}$ and the neighboring genes, *new25*$^{+}$ and *tam14*$^{+}$, are indicated by arrows below each panel. Note that *ade6*$^{+}$x8 *epe1Δ* R or W indicates whether clones analyzed exhibited Red (*ade6*-repressed) or White (*ade6*-expressing) phenotypes, respectively (see also Fig. 4E).

Unexpectedly, small RNA-seq analysis revealed that low but significant levels of small RNA are produced upstream of the *ade6*$^{+}$ TSS in some strains (upper and middle). These small RNAs were detected in the *ade6*$^{+}$x1 strain (*ade6*$^{+}$x1 *epe1Δ*), suggesting that their production is not due to *ade6*$^{+}$ repetition. Furthermore, these small RNAs were detected even in the absence of *ade6* siRNAs (*ade6*$^{+}$x8 *epe1Δ* R *ade6*-hp II*Δ*), indicating that the mechanism underlying the production of small RNAs produced upstream of the *ade6*$^{+}$ TSS is independent of, and different to, the mechanism responsible for production of *ade6*$^{+}$ siRNAs. Nonetheless, the RNAi pathway is responsible for the production of these small RNAs because they were abolished by loss of Dicer (*ade6*$^{+}$x8 *dcr1Δ ade6*-hp II*Δ*). Notably, all strains producing these small RNAs upstream of the *ade6*$^{+}$ TSS had high levels of H3K9me in this region (lower). Because H3K9me itself promotes the localization of the RDRC (Hayashi et al. 2012; Rougemaille et al. 2012), these results suggest that transcription coming from the outside region to robust ectopic heterochromatin results in recognition by the RDRC, followed by the production of small RNAs. Indeed, while a significant overlap between these small RNAs and H3K9me was observed upstream of the *ade6*$^{+}$ TSS, an adjacent *new25*$^{+}$, a possible source of such transcription, was not covered by H3K9me in these strains. Notably, the presence and absence of these siRNAs correlated with red (*ade6*-repressed) and white (*ade6*-expressing) phenotypes of *ade6*$^{+}$x8 *epe1Δ* strains (*ade6*$^{+}$x8 *epe1Δ ade6*-hp II*Δ* cells R or W), although their H3K9me levels on *ade6*$^{+}$ were comparable. The *ade6*$^{+}$x8 strains formed white (*ade6*-expressing) colonies even when only one copy of *ade6*$^{+}$ was derepressed by the loss of H3K9me. On the other hand, the H3K9me domain of *ade6*$^{+}$x8 *epe1Δ* W cells was comparable to that of R cells in ChIP-seq analysis because H3K9me levels at each *ade6*$^{+}$ copy cannot be determined in *ade6*$^{+}$x8 allele. Therefore, this correlation between the white (*ade6*-expressing) phenotype and the disappearance of small RNAs may support the hypothesis that the status of H3K9me itself affects the production of small RNAs upstream of the *ade6*$^{+}$ TSS. We assume that *ade6*$^{+}$x8 *epe1Δ* W cells lost H3K9me at the outermost copy of the *ade6*$^{+}$ repeats that will accept transcription from an outside region, therefore resulting in failure to produce these small RNAs.
