## Supplemental methods for "Tandemly repeated genes promote RNAi-mediated heterochromatin formation via an anti-silencing factor Epe1 in fission yeast"

<u>Generation of *ade6*$^+$ repeat strains</u>

Primers used for construction of *ade6*$^+$ repeat strains are listed in Supplemental Table S2. To generate repeated *ade6*$^+$ DNA fragments targeted to the endogenous *ura4*$^+$ locus, the *ade6*$^+$ gene fragment was duplicated stepwise on the plasmid, in which amplified fragments are flanked by the 5' and 3' end region of *ura4*$^+$. The backbone plasmid pAP-339 was first created as follows. PCR products of both upstream and downstream region of *ura4*$^+$ locus, and of the neomycine resistance gene (neo)-hCVM promoter region derived from pTL2M5, were ligated by In-Fusion cloning (Takara, cat#Z9648N or Clontech, cat#639648) along with BbsI-Bst1107 fragments of pTL2M5 (FYP2047, provided by the National Bio-Resource Project (NBRP), Japan) to provide the origin of replication in *E.coli* and the ampicillin resistance gene. To remove the neomycine resistance gene (AG-714, 715) and unnecessary sequences (AG-720, 721), consecutive rounds of inverse PCR (iPCR) with divergent primers were performed using KOD Fx neo (TOYOBO, cat# KFX-201). Next, the hCMV promoter of pAP-339 was replaced with the *ade6*$^+$ gene fragment as follows: pAP-339 was digested by SpeI and NheI, and ligated with the *ade6*$^+$ PCR fragment (chromosome III, 1315461-1318504) using In-Fusion cloning. The unwanted SpeI site within the *ade6*$^+$ gene fragment was disrupted by site-directed mutagenesis using iPCR (AG-751,752). After each step of iPCR or subcloning, the plasmids were sequenced. The resulting *ade6*$^+$x1 plasmid, pAP-444, has a PvuI site located in the middle of the ampicillin resistance gene, an SpeI site at the 5' end of the *ade6*$^+$ fragment, and an NheI site at the 3' end of the *ade6*$^+$ fragment (Supplemental Fig. S5A, left panel). Thus, when PvuI-*ade6*$^+$-SpeI and PvuI-*ade6*$^+$-NheI fragments derived from pAP-444 were ligated, the resultant plasmid had duplicate *ade6*$^+$ fragments, and the SpeI-NheI junction site became undigestible. Repetition of this duplication process resulted in *ade6*$^+$x2, x4, and x8 plasmids (pAP-464, pAP-485, pAP-523) (Supplemental Fig. S5A, middle panel). Furthermore, ligation of PvuI-*ade6*$^+$x2-SpeI (pAP-464) and PvuI-*ade6*$^+$x4-NheI (pAP-485) fragments resulted in the *ade6*$^+$x6 plasmid (pAP-599). These plasmids have NotI sites at either end of the *ura4*$^+$ fragment that allow retrieval of the *ade6*$^+$ repeat fragment targeting the endogenous *ura4*$^+$ locus. To promote homologous recombination, the CRISPR/Cas9 system was used to induce double strand breaks (DSB) at the endogenous *ura4*$^+$ locus (Supplemental Fig. S5A, right panel). Cloning of a gRNA sequence targeting the *ura4*$^+$ ORF into the gRNA-Cas9 single plasmid, pAH237 (Hayashi and Tanaka 2019), was performed as previously described, resulting in pAP-562. Co-transformation of the *ade6*$^+$ repeat fragment and gRNA-Cas9 plasmids targeting *ura4*$^+$ was carried out using the classic LiOAc method; $1 \times 10^8$ cells were transformed with 1 μg of AP-562 and 500–1000 ng of *ade6*$^+$ repeat fragments. To obtain clones in which the endogenous *ura4*$^+$ gene is replaced by the *ade6*$^+$ repeat, the transformants selected by EMM-selective plates (-leucine) were subsequently replicated to 5-FOA plates. The cells were again streaked on 5-FOA plates to obtain single colonies, and colony PCR was performed to select candidates. To isolate cells that had lost the gRNA-Cas9 plasmid, single colonies of each candidate were repeatedly streaked on YES plates until cells exhibited leucine auxotrophy on EMM (-leu) plates. Finally, genomic DNA was extracted from each candidate, and subjected to qPCR and Southern blot analysis to select clones in which the endogenous *ura4*$^+$ gene had been replaced by the *ade6*$^+$ repeat as planned (Supplemental Fig. S5B, C). Correlation between the copy number of the *ade6*$^+$ gene and the expression level of *ade6*$^+$ mRNA was confirmed by Northern blot and qRT-PCR (Supplemental Fig. S5D-F).

Generation of hairpin RNA constructs

Primers used for generation of *ade6* hairpin constructs are listed in Supplemental Table S2. The structure of the plasmid used to generate cells expressing *ade6* hairpin RNA is shown in Supplemental Fig. S6A. To generate an *ade6* hairpin construct whose expression is driven by the thiamine-repressible *nmt1*$^{+}$ promoter, the pREP1 plasmid (Maundrell 1993) was used as a backbone. The *ade6* hairpin construct was composed of three fragments. The left and right arms were PCR products corresponding to part of the *ade6*$^{+}$ ORF (*ade6* hairpin I, 250 bp: *ade6* hairpin II, 750 bp). For the third fragment, PCR products of the first intron of the *S.pombe rad9*$^{+}$ gene were used as a spacer. Using In-Fusion cloning (Takara, cat#Z9648N or Clontech, cat#639648), these three fragments were assembled and cloned in the pREP1 plasmid, which had first been linearized by SmaI and NdeI. To integrate this hairpin RNA construct at the endogenous *leu1*$^{+}$ locus, the *leu1*$^{+}$ targeting plasmid, pAP-509, was created, in which PCR products encoding the *leu1*$^{+}$ gene were subcloned into the SpeI-NotI site of pBluescript KS II (+). The pREP1-*ade6* hairpin construct was digested with PstI and EcoRI to retrieve fragments encoding the *nmt1*$^{+}$ promoter -*ade6* hairpin -*nmt1*$^{+}$ terminator fragment. These fragments were inserted into PstI-EcoRI digested pAP-509, creating the plasmids pAP-512 (*ade6*-hairpin I) and pAP-513 (*ade6*-hairpin II) (Supplemental Fig. S6A). To integrate these plasmids at the endogenous *leu1*$^{+}$ locus, the NruI site within the *leu1*$^{+}$ gene fragment was used for linearization, and the resultant fragments were transformed into *leu1-32* strains. Transformants were selected for leucine prototrophy, and integration at *leu1-32* was verified by colony PCR and subsequent Southern blot analysis. Strains carrying the *ade6* hairpin construct were created in the presence of thiamine to repress hairpin RNA expression during manipulation. To induce expression of hairpin RNA, cells were grown in liquid media without thiamine for 16 h and spread onto PMG plates without thiamine. Resultant colonies were stocked for use in experiments. Generation of *ade6* small RNAs in the various mutants was verified by Northern blot analysis (Supplemental Fig. S6B).

Southern blot analysis

Genomic DNA extraction was performed as described previously (https://www.baumann--lab.org/documents/Nurselab_fissionyeasthandbook_000.pdf). Genomic DNA was digested overnight with the indicated restriction enzymes and separated by agarose gel electrophoresis using 1xTris-Acetate-EDTA buffer, and then transferred to Amersham Hybond-N+ membranes (GE Healthcare, cat# RPN303B) using the TurboBlotter System (Cytiva, cat#10416328). The blots were UV cross-linked using a UVP CL-1000 Ultraviolet Crosslinker (120,000 $\mu J/cm^2$). The DNA probes, which were labeled with [$\gamma$-$^{32}$P] ATP by T4 polynucleotide kinase (Takara, cat# 2021A), were hybridized to the membrane in PerfectHyb Plus Hybridization buffer (Sigma, cat# H7033) at 42℃ overnight. The membrane was subsequently washed with 2× SSC 0.1% SDS buffer at 42℃. The imaging plate was exposed to the membrane for 1–2 days. The primers used for this analysis are listed in Supplemental Table S2.

Northern blot analysis

Total RNA was prepared using the hot phenol procedure and treated with recombinant DNase I (Takara, cat# 2270A) in the presence of RNasin Plus (Promega, cat# N2611). Samples were subjected to electrophoresis in 1% agarose with 6.7% formaldehyde and transferred to Amersham Hybond-N+ membranes (GE Healthcare, cat# RPN303B) using the TurboBlotter System (Cytiva, cat#10416328). Hybridization was performed as for Southern blot analysis. The primers used for this analysis are listed in Supplemental Table S2.

Additional analysis of small RNA-seq

For the truncation analyses of *cenH* (Supplemental Fig. S4B), only uniquely mapped reads were used with the -m 1 parameter. The Bedgraph files for visualizing in the IGV were produced with the "genomecov" function of the BEDTools program (version 2.26), which were then normalized by 1 million total mapped reads. For categorization of smRNA reads according to the annotation (Supplemental Fig. S8A), the "coverage" function of the BEDTools program was used. Analyses of smRNA length (Supplemental Fig. S8B,C) were conducted after extracting the reads mapped to the *ade6*$^{+}$x1 construct or pericentromeric *dg*/*dh* regions using the "intersect" function of the BEDTools program. The length information of each read is based on bed file. The 5'-end nucleotide analyses (Supplemental Fig. S8D,E) were performed using the WebLogo program (Crooks et al. 2004).

5'/3'RACE analysis of *cenH* and *Kint2::ura4*$^{+}$

Total RNA was extracted using the hot phenol method from 700 × $10^6$ cells, which were derived from exponentially growing cultures. This total RNA was treated with recombinant DNase I (Takara, cat# 2270A) in the presence of RNasin Plus (Promega, cat# N2611), and PolyA+ RNA was purified using the polyA Tract mRNA isolation system IV (Promega, cat# Z5310) according to the manufacturer's instructions. 5'/3'RACE analysis was performed with the SMARTer RACE 5'/3'Kit (Clontech, cat# 634858) following the manufacturer's instructions. The SMART technology takes advantage of a property of terminal transferase activity of reverse transcriptase, called template switching, to select 5'-ends of capped transcripts (Plessy et al. 2010). Instead of the provided SeqAmpDNA polymerase and linearized pRACE vector, KOD Fx neo (TOYOBO, cat# KFX-201) and a homemade vector derived from pUC18 were used for the amplification and cloning steps, respectively. To enhance specificity, nested PCR was used for analysis of *cenH* ncRNAs. The first PCR of 5'-RACE was performed with 45 cycles at 94℃ for 30 s and 68℃ for 3 min. The subsequent second PCR was performed using 5 µL of 50-fold diluted first-PCR products with 25 cycles at 94℃ for 30 s and 68℃ for 3 min. The first PCR of 3'-RACE was performed with 5 cycles at 94℃ for 30 s and 72℃ for 3 min, followed by 5 cycles at 94℃ for 30 s and 70℃ for 3 min, and finally 25 cycles at 94℃ for 30 s and 68℃ for 3 min. The second PCR was performed using 5 µL of 50-fold diluted first-PCR products with 25 cycles at 94℃ for 30 s and 68℃ for 3 min. For RACE analysis of *Kint2::ura4*$^{+}$ and *act1*$^{+}$, PCR was performed in the same way as for the first PCR of *cenH*. The results of sequencing RACE products, which were randomly selected from independent clones, were verified by BLAST, and it was confirmed that they were derived from the *cenH* element, not from the pericentromeric *dg*/*dh* element. For sense transcripts and antisense transcripts of *cenH* ncRNA, 30 clones of 5'RACE or 3'RACE products were identified. The primers used for this analysis are listed in Supplemental Table S2. The sequence data of RACE products are listed in Supplemental Data S1.

Truncation analysis of *cenH* fragment with a plasmid-based minichromosome

To generate truncated *cenH* fragments, the 4.5 kb *cenH* sequence was amplified by PCR from genomic DNA of the $h^{90}$ strain. Diluted PCR products were used as template for the subsequent PCRs, which produced a series of truncated *cenH* fragments. The resultant PCR products had BamHI and NcoI sites derived from primers and were cloned into the corresponding site of MC-L5 (Buscaino et al. 2013). This resulted in a plasmid-based minichromosome in which the L5 fragment is replaced by a truncated *cenH* fragment (pAP-275, pLCC-*cenH* 3.0 kb; pAP-315, pLCC-*cenH majorΔ Rev TSS*; pAP-273, pLCC-*cenH major Rev TSS only*; pAP-269, pLCC-*cenH* 1.5 kb). Strains containing minichromosomes were grown in PMG medium lacking adenine and uracil. Primers used for generation of the pLCC-*cenH* series are listed in Supplementary Table S2.

Additional analyses of CAGE-seq

For consensus sequence analysis of core promoters in *dg*/*dh*, the sequences of ±7 nt or ±50 nt flanking unique CAGE TSSs (>0.05 CPM) were retrieved from both ±strands of *dg*/*dh* elements, specifically, chromosome I (3752407–3765008, 37772666–3789949), chromosome II (1602261–1618293, 1630123–1643849), chromosome III (1071176–1092437, 1106558–1139536), and MTR (1–20128), and analyzed by Weblogo3 (Crooks et al. 2004). As a control, the same number of pseudo TSSs was randomly chosen from anywhere in the *dg*/*dh* elements, and sequences of ±50 nt flanking those pseudo TSSs were analyzed in the same way. For analysis of mRNA TSSs, CAGE TSSs (>0.05 CPM) of the wild type mapped in the annotated 5'-UTR (including its upstream 100 nt) of euchromatic genes were classified into three groups (high, medium, and low) based on expression levels (CPM). For each group, the sequences of ±7 nt or ±50 nt flanking unique TSSs were retrieved and analyzed in the same way. To extract CAGE TSSs that had both the Inr (Y/R) sequence at -1/+1 and A/T rich region at 25–32 nt upstream of the TSS, a Position-Weight matrices file was created by Biopython (https://biopython.org/) with FASTA files that were used for Weblog analysis. The resultant matrices file was consistent with that of WebLogo. Based on this matrices file, CAGE TSSs with both the Inr sequence and the upstream A/T rich region were extracted using FIMO (http://meme-suite.org/doc/fimo.html?man_type=web) and mapped to ASM294v2 genomes.
